# NIPBL and WAPL balance cohesin activity to regulate chromatin folding and gene expression

**DOI:** 10.1101/2022.04.19.488785

**Authors:** Jennifer M. Luppino, Andrew Field, Son C. Nguyen, Daniel S. Park, Parisha P. Shah, Yemin Lan, Rebecca Yunker, Rajan Jain, Karen Adelman, Eric F. Joyce

## Abstract

The relationship between cohesin-mediated chromatin looping and gene expression remains unclear. We investigated the roles of NIPBL and WAPL, two regulators of cohesin activity, in chromatin folding and transcription in human cells. Consistent with their opposing roles in cohesin regulation, depletion of these factors showed opposite effects on levels of chromatin-bound cohesin and spatial insulation of neighboring domains. We find that NIPBL or WAPL depletion each alter the expression of ~2,000 genes, most of which are uniquely sensitive to either regulator. We find that each set of differentially expressed genes are enriched at chromatin loop anchors and clustered within the genome, suggesting there are genomic regions sensitive to either more or less cohesin. Remarkably, co-depletion of both regulators rescued chromatin misfolding and gene misexpression compared to either single knockdown. Taken together, we present a model in which the relative, rather than absolute, levels of NIPBL and WAPL are required to balance cohesin activity in chromatin folding to regulate transcription.

## Introduction

The concept that structure informs function is fundamental to many aspects of biology, yet the direct relationship between the three-dimensional folding of the genome and gene expression remains unclear. Chromosome conformation capture-based techniques have revealed a complex hierarchy of structural layers that help to organize mammalian chromosomes (*1–4*). In particular, chromosomes are segmented at the sub-megabase scale into topologically associating domains (TADs) (*5–8*), which are well conserved across cell types and even species (*9*).

TADs are believed to create a favorable environment for transcription by facilitating communication between gene promoters and their regulatory elements (*10–16*). The prevailing hypothesis is that TADs form via loop extrusion, a process in which the highly conserved ring-shaped cohesin complex actively compacts chromatin until it encounters convergently oriented CTCF sites (*17–21*). The frequent stalling of cohesin at a CTCF-binding site tends to define the boundaries of TADs and anchors of loops. However, live and fixed imaging of loops and TADs in single cells has revealed that these structures are extremely dynamic and heterogenous across a cell population (*22–27*).

The dynamics underlying cohesin-mediated chromatin looping depends on the interplay between two mutually exclusive regulators of cohesin, Nipped-B-like protein (NIPBL) and Wings apart-like protein homolog (WAPL) (*28–30*). NIPBL has been proposed to both load cohesin and activate its ATPase domain to initiate loop extrusion (*31, 32, 30, 33, 34*). In contrast, WAPL removes cohesin from chromatin, limiting its residence time to minutes, thus restricting the size of loops and TADs across the genome (*35–37, 29, 18, 38*). While complete or near complete loss of cohesin leads to widespread, albeit modest, effects on gene expression (*21, 38–40*), it remains unclear how perturbation of NIPBL or WAPL across multiple cell divisions would influence gene regulation.

In this study, we aimed to understand this question by depleting NIPBL or WAPL from chromatin across several cell cycles. We find that ~90% loss of NIPBL alters the spatial insulation between TADs to a similar extent as complete cohesin loss but without affecting mitosis or cell growth. Interestingly, we find that NIPBL or WAPL depletion leads to misexpression of unique sets of genes; nonetheless, these genes have many shared features, including proximity to loop anchors, cohesin, and each other. Indeed, differentially expressed genes are clustered within domains and exhibit coordinated misexpression, suggesting there are differential genomic regions with increased sensitivity to altered levels of cohesin. Remarkably, co-depletion of both regulators rescued the spatial insulation between TADs and gene misexpression compared to either single knockdown. We propose that a stoichiometric balance between NIPBL and WAPL may be essential for normal cohesin function. Together, these studies provide insights into how cohesin is dynamically regulated by opposing cofactors to organize chromatin and facilitate proper gene regulation.

## Results

### Depletion of NIPBL and WAPL influence chromatin folding in a locus-specific manner

To determine the effects of NIPBL or WAPL depletion through multiple cell divisions, we optimized a 72-hour small interfering RNA (siRNA) protocol in human HCT-116 cells (**Fig. 1A, and fig. S1, A and B**). We performed subcellular protein fractionation followed by quantitative western blotting with fluorescence detection and found a robust 92% and 89% reduction in chromatin-bound levels of NIPBL and WAPL, respectively (**Fig. 1, B and C**). siNIPBL resulted in a 38% depletion of chromatin-bound RAD21 levels whereas siWAPL increased chromatin-bound RAD21 levels by 18% (**Fig. 1, D and E**). NIPBL or WAPL depletion did not change the growth rate of cells over the course of four cell divisions and did not alter mitotic progression, chromosome segregation or the frequency of mitotic entry (**Fig. 1, F to H, and fig. S1, C to D**).

**Figure. 1:**
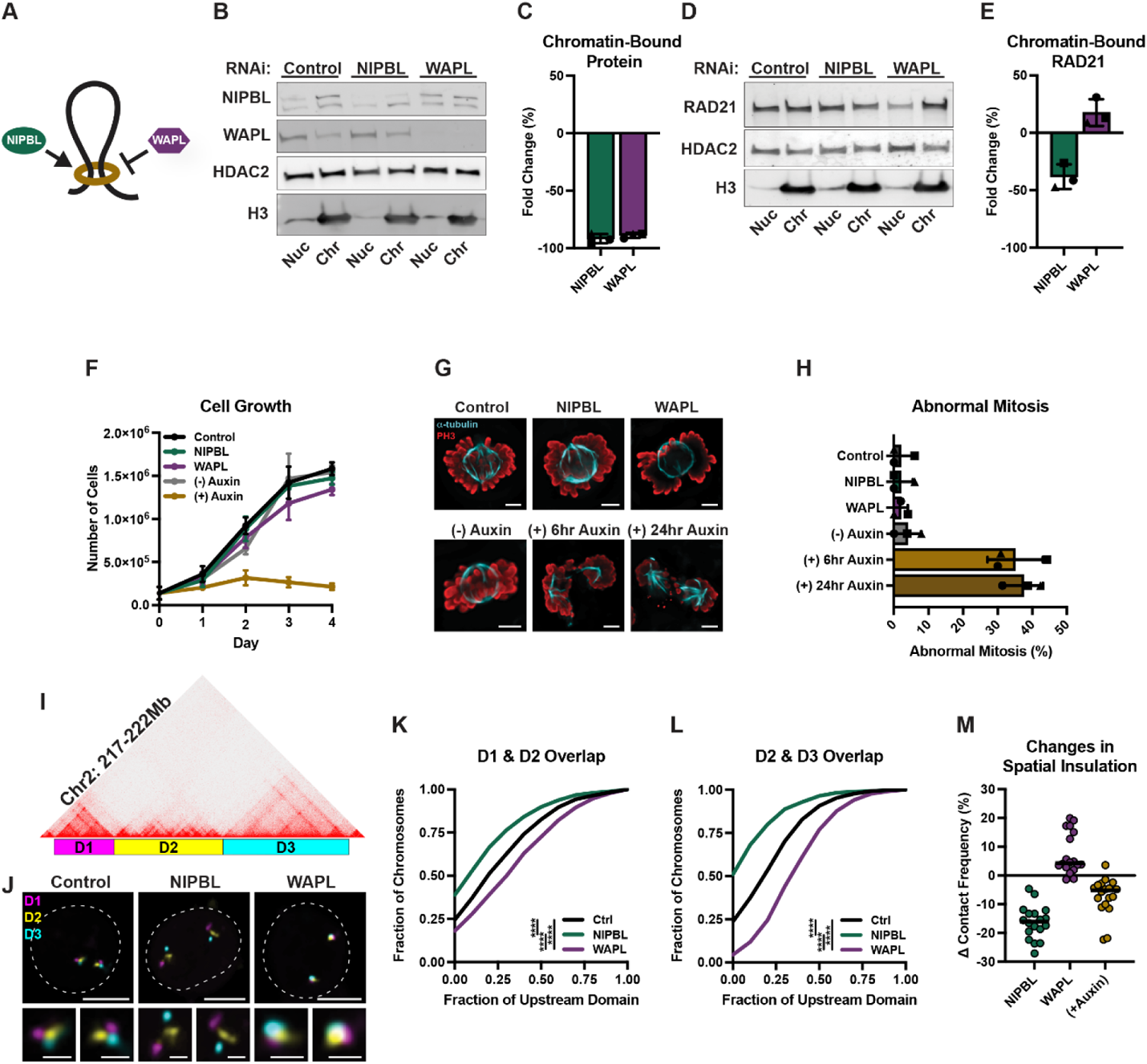
Depletion of NIPBL and WAPL influence chromatin folding in a locus-specific manner. (**A**) Cartoon depicting the roles of the two opposing cohesin regulators; NIPBL loads cohesin onto chromatin and is required for loop extrusion whereas WAPL opens the ring and removes it. (**B**) Fluorescent western blot to NIPBL (top of the two bands) and WAPL in nuclear (nuc) and chromatin-bound (chr) subcellular protein fractionations of RNAi control, NIPBL, or WAPL depleted HCT-116 cells. All bands are from the same blot. (**C**) Mean fold change (%) of NIPBL and WAPL bound to chromatin in each respective knockdown. Each symbol represents a biological replicate, error bars represent standard deviation. (**D**) Fluorescent western blot to RAD21 in nuclear (nuc) and chromatin-bound (chr) subcellular protein fractionations of RNAi control, NIPBL, or WAPL depleted HCT-116 cells. All bands are from the same blot. (**E**) Mean fold change (%) of RAD21 bound to chromatin in each respective knockdown. Each symbol represents a biological replicate, error bars represent standard deviation. (**F**) Cell growth measured in 24-hour increments following RNAi or auxin treatment. Each bar represents the mean of 3 biological replicates and error bars represent the standard deviation. (**G**) Representative immunofluorescence images of mitotic cells stained for α-tubulin (cyan) and phospho-Histone H3 (PH3; red) in RNAi control, NIPBL, or WAPL depleted HCT-116 cells and HCT-116-RAD21-AID cells −/+ auxin for 6 or 24 hours. Scale bar, 5μm. (**H**) Average percentage of abnormal mitotic cells in RNAi control, NIPBL, or WAPL depleted HCT-116 cells and HCT-116-RAD21-AID cells −/+ auxin for 6 or 24 hours. Each symbol represents a biological replicate, error bars represent standard deviation. (**I**) Oligopaint design for three neighboring domains at chr2:217-222Mb. (**J**) Representative FISH images for three domains at chr2:217-222Mb in RNAi control, NIPBL, and WAPL depleted HCT-116 cells. Dashed line represents nuclear edge, scale bar, 5μm (above) or 1μm (below). (**K**) Cumulative frequency distribution of overlap between the neighboring domains D1 and D2 on chr2 in RNAi control (n = 1,170 chromosomes), NIPBL (n = 1,177 chromosomes), or WAPL (n = 1,136 chromosomes) depleted HCT-116 cells. Two-tailed Mann-Whitney test, **** p < 0.0001. (**L**) Cumulative frequency distribution of overlap between the neighboring domains D2 and D3 on chr2 in RNAi control (n = 1,202 chromosomes), NIPBL (n = 1,284 chromosomes), or WAPL (n = 1,149 chromosomes) depleted HCT-116 cells. Two-tailed Mann-Whitney test, **** p < 0.0001. (**M**) Change in contact frequency across 18 domain pairs in NIPBL, or WAPL depleted HCT-116 cells and HCT-116-RAD21-AID cells treated with auxin for 6 hours. Each dot represents the median of ≥ 4 biological replicates at each locus.

In contrast to NIPBL or WAPL depletion, near-complete loss of RAD21 (<10% remaining on chromatin; **fig. S1, E and F**) via auxin-inducible degradation (AID) prompted growth arrest after the first cell division (**Fig. 1F**) and resulted in a higher mitotic index after only 6 hours of auxin treatment (**fig. S1C**). Most mitotic cells were arrested in metaphase (**fig. S1D**) with morphological abnormalities, including precocious anaphase, multi-polar spindles, chromosome congregation defects, and lagging chromosomes (**Fig. 1, G and H**). We also observed an increase in the frequency of nuclear blebbing and micronuclei, consistent with chromosome segregation errors. These results confirm that cohesin is essential for normal mitotic progression in HCT-116 cells. These data further suggest that cells do not require the full complement of NIPBL or WAPL for cell growth or fidelity. Thus, a small amount of NIPBL across several cell divisions is sufficient to load most cohesin onto chromatin.

Next, to determine whether decreased NIPBL or WAPL might be sufficient to alter chromatin folding, we used an Oligopaint fluorescence *in situ* hybridization (FISH)-based assay that we previously developed to quantify the frequency of interactions across domain boundaries as measured by the extent of spatial overlap (*25*). We labeled three consecutive domains on chromosome 2 that had strong intervening boundaries (20th and 6th percentiles, as measured by Hi-C insulation scores) (**Fig. 1I**). Neighboring domains exhibited less spatial overlap in cells depleted of NIPBL than in control cells, consistent with chromatin misfolding of the labelled locus (**Fig. 1, J to L**). The extent of spatial separation was similar to that observed after acute (6 hour) and near complete degradation of RAD21 (**fig. S1, G to I**). In contrast, WAPL depletion led to increased interactions across both domain boundaries (**Fig. 1, J to L**).

We expanded our FISH assay to label sixteen additional domain or sub-domain boundaries (**fig. S1J**). The Oligopaint probes spanned regions of different boundary strengths (defined by their insulation score), gene densities, and chromatin types. We used a recently developed high-throughput FISH platform, called HiDRO, to image at least four biological replicates of each FISH reaction in parallel (*41*). We defined a contact cutoff of 250 nm based on the resolution of our microscope to quantify interactions across domain boundaries. As expected, cohesin loss by siNIPBL or RAD21 AID decreased the contact frequencies across all boundaries assayed with variable locus sensitivities (**Fig. 1M, and fig. S1, K to M**). We observed a 5–28% and 2–22% decrease in contact frequency after NIPBL depletion and RAD21 AID, respectively (**Fig. 1M, and fig. S1M**). WAPL depletion increased contact across most boundaries, with a 1– 20% increase in contact frequency in 16/18 domain pairs (**Fig. 1M, and fig. S1M**). Therefore, ~90% loss of either protein was sufficient to alter chromatin folding by FISH but did not affect cell growth or proliferation, suggesting that a small amount of NIPBL and WAPL is sufficient for proper sister-chromatin cohesion and chromosome segregation.

### NIPBL and WAPL regulate the expression of different genes

We next sought to determine the extent of gene expression changes after siRNA depletion of NIPBL or WAPL. We performed precision nuclear run-on sequencing (PRO-seq) to map the locations of active RNA polymerases and to determine levels of nascent transcription across two biological replicates. Given the reproducibility between our replicates (Spearman’s rho ≥0.95), we merged the data within each condition for downstream analyses.

To define differentially expressed genes (DEGs), we applied the DESeq2 algorithm and further filtered significant DEGs for a minimum adjusted p-value of 0.01. We identified 1,877 and 1,932 DEGs after NIPBL or WAPL depletion, respectively (**Fig. 2, A and B**). Most changes were modest, and >95% of the DEGs had less than a two-fold change in expression (**Fig. 2, A and B**). Genes were approximately equally up- and downregulated in each knockdown condition (53% upregulated and 47% downregulated DEGs after NIPBL knockdown; 47% upregulated and 53% downregulated DEGs after WAPL knockdown). These results resemble that of acute RAD21 depletion in the same cell line (*40*). This suggests that NIPBL and WAPL modify both chromatin folding and gene expression to a similar extent as acute RAD21 degradation despite differences in cell survival outcomes.

**Figure 2:**
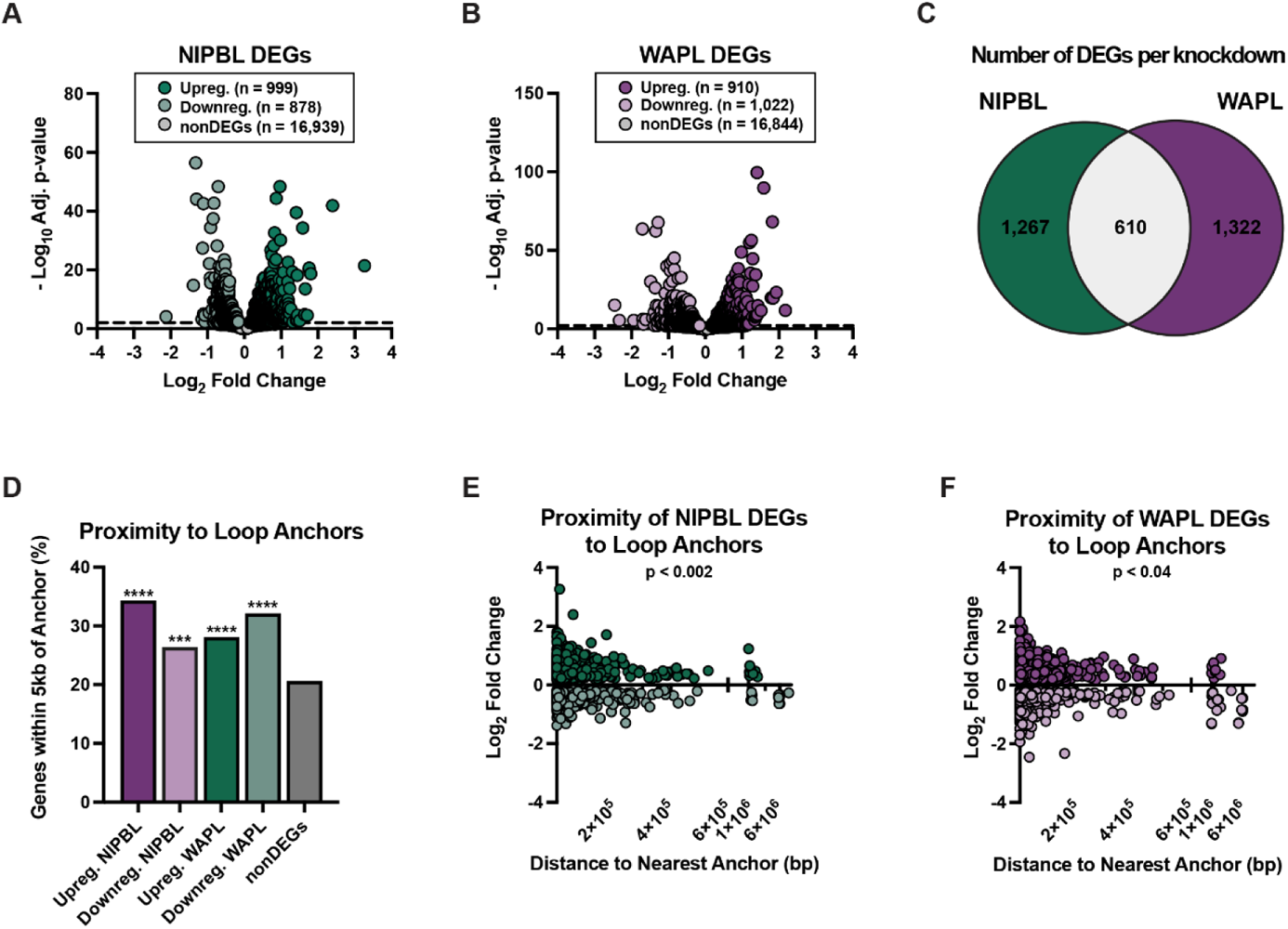
NIPBL and WAPL regulate the expression of different genes. (**A**) The log_2_(fold change) of genes after NIPBL knockdown versus their significance. DEGs are in green (999 up, 878 down) and non-significantly changed genes (nonDEGs, adjusted p-value > 0.01) are in grey. (**B**) The log_2_(fold change) of genes after WAPL knockdown versus their significance. DEGs are in purple (910 up, 1022 down) and nonDEGs (adjusted p-value > 0.01) are in grey. (**C**) Venn diagram of the number of NIPBL, WAPL, and shared DEGs. (**D**) Percentage of up, down, NIPBL, WAPL, or nonDEGs with a TSS within 5kb of a loop anchor. Fisher’s exact test compared to nonDEGs, **** p < 0.0001, *** p = 0.0002. (**E**) Distance from each NIPBL DEG TSS to the nearest loop anchor versus the fold change of the gene. Spearman correlation, p = 0.0018. (**F**) Distance from each WAPL DEG TSS to the nearest loop anchor versus the fold change of the gene. Spearman correlation, p = 0.037.

Genes differentially expressed after RAD21 and NIPBL depletion were enriched in the same top four Gene Ontology (GO) terms for biological processes (**fig. S2, A and B, and Tables S1 and S2**). However, when comparing NIPBL to WAPL depletion, >70% of the DEGs were unique to either condition (**Fig. 2C**). Moreover, the top GO terms differed between NIPBL- and WAPL-sensitive genes, indicating that not only different genes but also different pathways were predominantly affected by the two knockdowns (**fig. S2, A and C, and Tables S1 and S3**). This suggests that a minority of sites are equally sensitive to both NIPBL and WAPL depletion. Surprisingly, of the relatively few shared DEGs (610 genes) between NIPBL and WAPL, >80% (473 genes) were changed in the same direction, with equal rates of up- and downregulation observed (**fig. S2D**).

Despite their differences, the unique NIPBL- and WAPL-sensitive genes shared many features. When compared with the position of chromatin loops from our Hi-C dataset, we found that 95% of the DEGs in either condition were within 200 kb of a loop anchor and ~30% were within 5 kb. In comparison, only 20% of the nonDEGs were found within 5kb of a loop anchor (**Fig. 2D**). Moreover, we found that genes closer to anchors tended to have a greater fold change in expression (**Fig. 2, E and F**). This is similar to our previous observations following acute loss of RAD21 (*25*), highlighting a general signature of cohesin dysfunction in which genes at loop anchors are especially sensitive.

As chromatin loops are typically enriched for both cohesin and the insulator binding protein CTCF, we performed chromatin immunoprecipitation sequencing (ChIP-Seq) across four biological replicates to map their co-localization genome-wide. The promoters of DEGs were significantly enriched for RAD21 and CTCF co-occupancy compared with the promoters of nonDEGs (**fig. S2E**). Taken together, these data indicate that genes sensitive to NIPBL or WAPL depletion are predominantly unique to either condition but are both found near structural features and bound by architectural proteins.

### Cohesin-sensitive genes are clustered and coordinated within TADs

To further investigate the relationship between gene expression and chromatin topology, we asked whether genes differentially expressed after NIPBL or WAPL knockdown were arranged randomly throughout the genome or instead clustered within TADs. Active genes were assigned to one of 3,342 TADs across the genome, with each TAD harboring an average of 32 genes. To determine if DEGs sensitive to NIPBL or WAPL depletion clustered significantly more than expected, we computationally permutated the assignment of DEG or nonDEG to all active genes 1,000 times to create null distributions for each knockdown condition (**Fig. 3, A and B**). We found that both NIPBL- and WAPL-sensitive genes were clustered in TADs significantly higher than expected by chance (**Fig. 3, A and B**).

**Figure 3:**
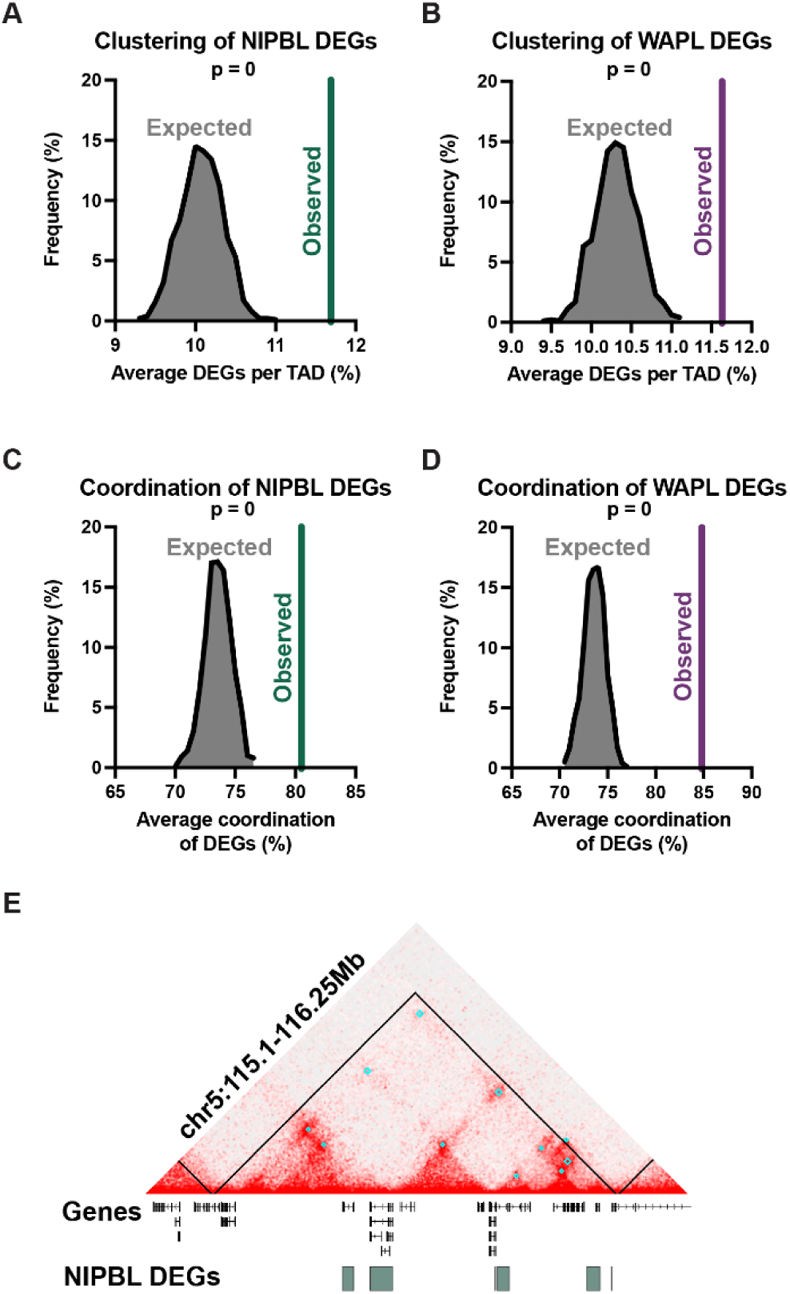
Cohesin-sensitive genes are clustered and coordinated within TADs. (**A**) The observed average percentage of NIPBL DEGs per TAD compared to a null distribution (expected). Permutations generated by shuffling the DEG and nonDEG designations across genes 1,000 times. Analysis limited to TADs with at least one expressed gene. Exact test, p = 0. (**B**) The observed average percentage of WAPL DEGs per TAD compared to a null distribution (expected). Permutations generated by shuffling the DEG and nonDEG designations across genes 1,000 times. Analysis limited to TADs with at least one expressed gene. Exact test, p = 0. (**C**) The average coordination of NIPBL DEGs compared to a null distribution generated by shuffling the fold change amongst the DEGs 1,000 times. Analysis limited to TADs with at least two expressed genes. Exact test, p = 0. (**D**) The average coordination of WAPL DEGs compared to a null distribution generated by shuffling the fold change amongst the DEGs 1,000 times. Analysis limited to TADs with at least two expressed genes. Exact test, p = 0. (**E**) Representative TAD with 100% DEG coordination on chr5 which contains six downregulated NIPBL DEGs. Black lines represent TADs, cyan boxes represent loops.

We then investigated whether DEGs in each TAD have coordinated changes in their expression. For each TAD containing at least two DEGs, we calculated a coordination score based on the directionality of gene expression changes. Random expression would yield 50% coordination within a TAD. In contrast, genes differentially expressed after NIPBL and WAPL knockdown were on average 80.5% and 84.8% coordinated, respectively, which was significantly greater than expected (**Fig. 3, C and D**). Moreover, we found that TADs with 90– 100% coordination were significantly enriched above the null distribution, whereas TADs with 50–60% coordination were significantly depleted (**fig. S3, A and B**). This suggests that DEGs are dysregulated in a coordinated fashion when they are found within the same TAD. Indeed, 52% and 60% of TADs with >2 DEGs had 100% coordination of genes differentially expressed after NIPBL and WAPL knockdown, respectively. This was especially apparent at a 1 Mb-sized TAD on the q arm of chromosome 5 that harbored six DEGs, all of which were downregulated after NIPBL knockdown (**Fig. 3E**). Similarly, a TAD on the q arm of chromosome 17 harbored seven DEGs, all of which were upregulated after NIPBL knockdown (**fig. S3C**). In both examples, the DEGs were also enriched at loop anchors.

Considering the high coordination of DEGs within TADs, we reasoned that enhancer(s) within a domain might be activated or repressed after knockdown, and therefore affect the expression of all nearby genes. Alternatively, changes in the spatial organization of chromatin within a TAD might elicit miscommunication between regulatory elements and promoters separate from altered enhancer activity. To distinguish between these possibilities, we identified putative enhancer elements from the PRO-seq data using the discriminative Regulatory-Element detection algorithm (dREG). dREG peaks were further refined to predict 23,741 active enhancers in HCT-116 cells. We then analyzed changes in PRO-seq signal at the dREG peaks to test whether eRNA synthesis, and thus enhancer activity, was changed in the knockdown conditions. We found that most (96%) enhancer peaks did not change after NIPBL or WAPL knockdown, suggesting that the changes in transcription were not caused by altered enhancer activity (**fig. S3, D and E**). Instead, these data along with our FISH results support a model in which changes in gene expression due to cohesin dysfunction are likely caused by changes in chromatin folding within TADs.

### Co-depletion of NIPBL and WAPL restores normal chromatin folding

The differential effects of NIPBL and WAPL depletion on both chromatin folding and gene expression motivated us to test whether they could possibly balance one another. We simultaneously knocked down each protein by 96% and 94%, respectively, similar to the single knockdown conditions (**Fig. 4, A and B**). Importantly, cell growth, mitotic entry, and chromosome segregation remained unaltered in the double knockdown condition, indicating that HCT-116 cells can tolerate simultaneous depletion of both proteins across a minimum of four divisions (**fig. S4, A to D**). We first measured cohesin levels after subcellular fractionation, which demonstrated a partial rescue of RAD21 levels on chromatin compared to the single knockdowns (**Fig 1D, and Fig. 4, C and D**). We next performed Oligopaint FISH to assess boundary strength in single cells as measured by the extent and frequency of spatial overlap between neighboring domains. Despite only partial rescue of chromatin-bound cohesin levels, the double knockdown restored the distribution of spatial overlap across two boundaries analyzed on chromosome 2 (**Fig. 4, E to G, and fig. S4, E and F**). Using HiDRO, we extended this assay to sixteen additional loci across the genome and found that all but one boundary showed partial or complete rescue of inter-domain interactions after double knockdown of NIPBL and WAPL (**Fig. 4H, and fig. S4G**).

**Figure 4:**
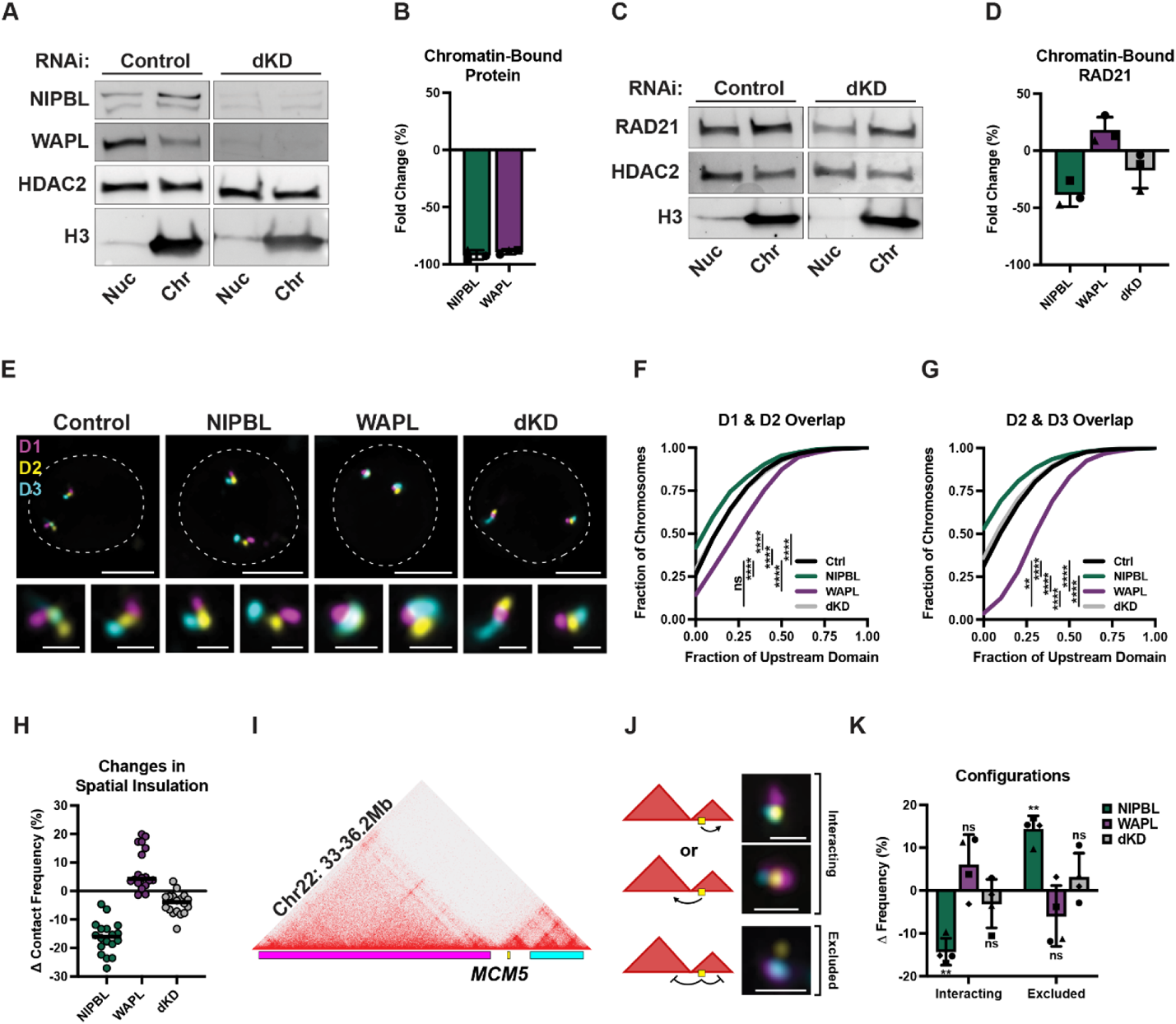
Co-depletion of NIPBL and WAPL restores normal chromatin folding. (**A**) Fluorescent western blot to NIPBL (top of the two bands) and WAPL in nuclear (nuc) and chromatin-bound (chr) subcellular protein fractionations of RNAi control and NIPBL and WAPL double knockdown (dKD) depleted HCT-116 cells. All bands are from the same blot. (**B**) Mean fold change (%) of NIPBL and WAPL bound to chromatin in the double knockdown condition. Each symbol represents a biological replicate, error bars represent standard deviation. (**C**) Fluorescent western blot to RAD21 in nuclear (nuc) and chromatin-bound (chr) subcellular protein fractionations of RNAi control and NIPBL and WAPL double knockdown (dKD) depleted HCT-116 cells. All bands from the same blot. (**D**) Mean fold change (%) of RAD21 bound to chromatin in RNAi control, NIPBL, WAPL, and double knockdown (dKD) depleted HCT-116 cells. Each symbol represents a biological replicate, error bars represent standard deviation. (**E**) Representative FISH images for three domains at chr2:217-222Mb in RNAi control, NIPBL, WAPL, and NIPBL and WAPL co-depleted HCT-116 cells. Dashed line represents nuclear edge, scale bar, 5μm (above) or 1μm (below). (**F**) Cumulative frequency distribution of overlap between the neighboring domains D1 and D2 on chr2 in RNAi control (n = 2,172 chromosomes), NIPBL (n = 1,514 chromosomes), WAPL (n = 1,704 chromosomes), or dKD (n = 1,620 chromosomes) depleted HCT-116 cells. Two-tailed Mann-Whitney test, **** p < 0.0001, ns = not significant (p = 0.79). (**G**) Cumulative frequency distribution of overlap between the neighboring domains D2 and D3 on chr2 in RNAi control (n = 2,188 chromosomes), NIPBL (n = 1,571 chromosomes), WAPL (n = 1,719 chromosomes), or dKD (n = 1,661 chromosomes) depleted HCT-116 cells. Two-tailed Mann-Whitney test, **** p < 0.0001, ** p = 0.0014. (**H**) Change in contact frequency across 18 domain pairs in HCT-116 cells depleted for NIPBL, WAPL, or both. Each dot represents the median of ≥ 4 biological replicates at each locus. (**I**) Oligopaint design to *MCM5* and neighboring domains at chr22:32-36.2Mb. (**J**) Cartoon diagrams and representative FISH images of the possible interactions between *MCM5* and its neighboring domains at chr22:32-36.2Mb. The “interacting” configuration is defined as the majority of the *MCM5* signal overlapping either the up or downstream domain. “Exclusion” is defined as the majority of *MCM5* signal non-overlapping with either neighboring domain. Dashed line represents nuclear edge, scale bar 1μm. (**K**) Change in the frequency of interacting and exclusion between *MCM5* and neighboring domains at chr22:32-36.2Mb in RNAi control, NIPBL, WAPL, or double knockdown cells. Each bar represents the mean of four biological replicates, error bars represent standard deviation. Two-tailed paired t-test, ** p = 0.003 interacting; p = 0.003 exclusion, ns = not significant (p ≥ 0.19).

Given the model that cohesin facilitates interactions between enhancers and promoters, we next examined whether the double knockdown might rescue interactions between a gene and its cis-regulatory domains. In this three-color FISH assay, the gene may interact with either of its neighboring domains or be excluded from both domains (**Fig. 4, I and J**). We focused on the boundary-proximal gene *MCM5,* which we found was displaced from its neighboring domains following acute degradation of RAD21 (**Fig. 4K**) (*25*). Again, co-depletion of NIPBL and WAPL restored the distribution of *MCM5*-domain configurations to that observed in the control samples (**Fig. 4K**). Together, these findings further support the notion that balancing the levels of NIPBL and WAPL can restore chromatin-bound cohesin levels and proper chromatin folding.

### NIPBL and WAPL balance cohesin activity to regulate gene expression

We next sought to determine whether rescue of chromatin folding by FISH was sufficient to normalize gene expression. We performed PRO-seq in cells co-depleted of NIPBL and WAPL and found approximately half as many significant DEGs (1,042 genes) in the double knockdown compared to either of the single knockdowns (**Fig. 5A**). These genes were approximately equally up- and downregulated (56% and 44%, respectively), similar to the results observed in the single knockdowns (**Fig. 2, A and B**). Most gene expression changes were modest, and 97% of DEGs showed less than two-fold change in expression (**Fig. 5A**).

**Figure 5:**
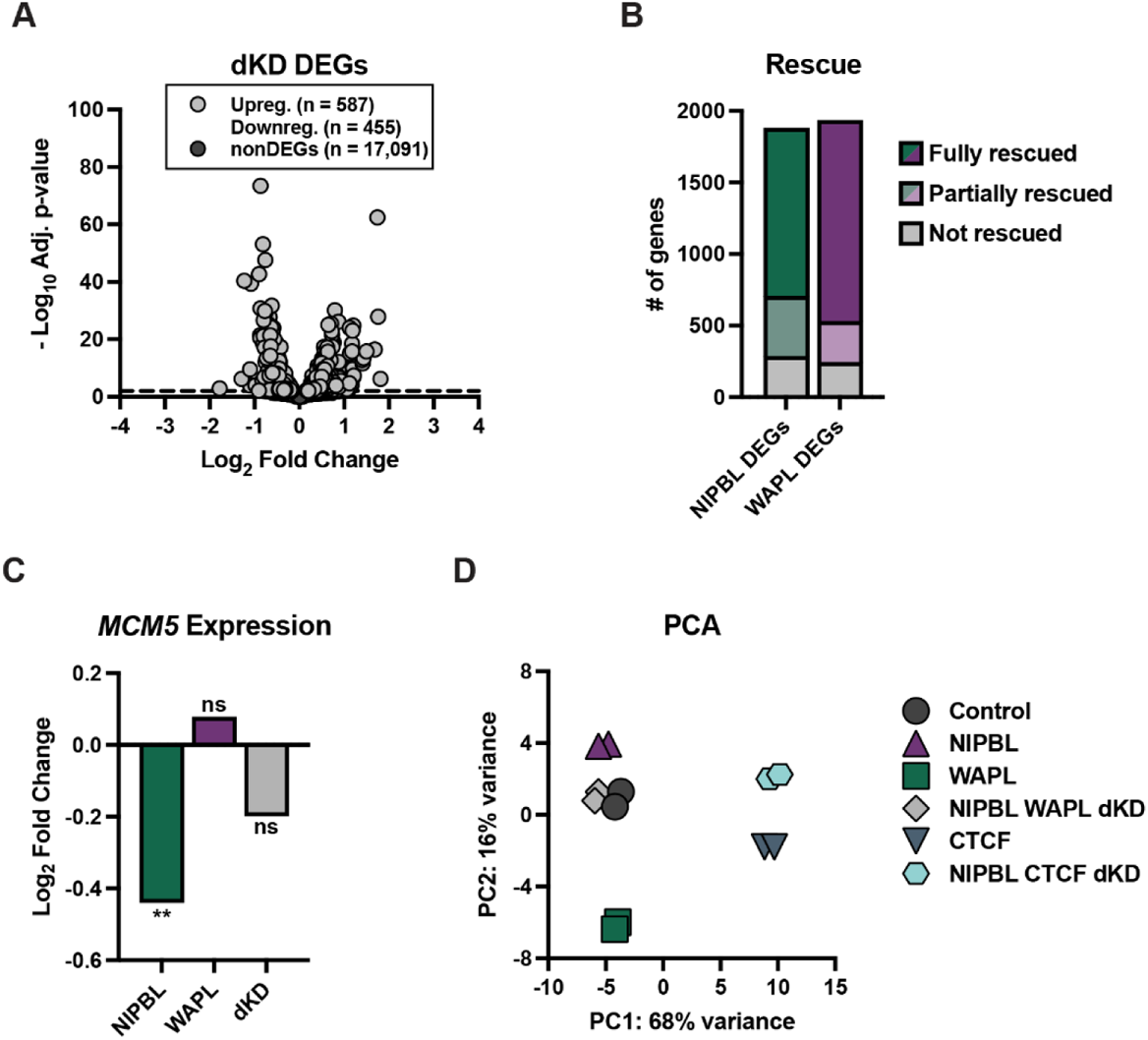
NIPBL and WAPL balance cohesin activity to regulate gene expression. (**A**) The log_2_(fold change) of genes after NIPBL and WAPL double knockdown versus their significance. DEGs are in light grey (587 up, 455 down) and non-significantly changed genes (adjusted p-value > 0.01) are in dark grey. (**B**) Number of NIPBL and WAPL DEGs fully, partially, or not rescued in the double knockdown condition. (**C**) The log_2_(fold change) of *MCM5* expression in the NIPBL, WAPL, or double knockdown conditions. ** p = 0.002, ns = not significant (p = 0.58 for WAPL, p = 0.13 for dKD). (**D**) Principal component analysis plot of the PRO-seq data. Each symbol represents one biological replicate of knockdown.

We compared the DEGs in the NIPBL knockdown and double knockdown conditions to determine which, if any, gene expression changes were rescued by co-depletion with WAPL. We found that 1,174 of the 1,877 DEGs identified in the NIPBL knockdown were completely rescued and were no longer significantly differentially expressed in the double knockdown (**Fig. 5B**). This included *MCM5,* which was downregulated by 1.5-fold after NIPBL knockdown but was no longer a significant DEG following WAPL co-depletion (**Fig. 5C**). Thus, restoring chromatin folding patterns at this locus was accompanied by a rescue of *MCM5* gene expression. Another 421 genes were partially rescued; these genes were still significantly misexpressed in the same direction as in the NIPBL single knockdown, but their fold change was diminished (**Fig. 5B**). Together, co-depletion of NIPBL and WAPL rescued 85% of the genes differentially expressed after NIPBL knockdown (**Fig. 5B**). These rescued genes were approximately equally up- and downregulated (841 upregulated and 754 downregulated) and were enriched in the same top six GO terms as the genes differentially expressed after NIPBL knockdown (**Tables S1 and S4**), suggesting that the major biological processes disrupted by NIPBL depletion can be rescued by double knockdown with WAPL.

We next reciprocally examined whether WAPL-dependent gene expression might also be rescued by co-depletion of NIPBL. Remarkably, of the 1,932 genes sensitive to WAPL depletion, 1,405 were fully rescued and another 287 were partially rescued in the double knockdown condition (**Fig. 5B**). In total, expression of 88% of genes differentially expressed after WAPL depletion was restored by co-depletion of NIPBL. Rescued DEGs represented both up- and down-regulated genes (769 and 923, respectively), and were enriched in similar biological processes to those genes differentially expressed after WAPL single knockdown (**Tables S3 and S5**). Taken together, these data show that co-depletion of NIPBL and WAPL can offset each other and correct for the majority of gene misexpression observed in either single depletion.

### CTCF loss partially rescues gene misexpression in NIPBL-depleted cells

Considering WAPL co-depletion with NIPBL could restore gene expression to normal levels, we next asked whether any opposing regulator of cohesin activity might have this capacity. Therefore, we next investigated whether co-depletion of CTCF, which inhibits loop extrusion by stabilizing cohesin on chromatin (*17, 20, 42*), might have a similar effect to that of WAPL depletion. CTCF knockdown alone significantly altered the expression of 3,889 genes (**fig. S5A**). As previously observed in other cell types, the majority of CTCF DEGs (92%) had modest fold changes (<two-fold change; **fig. S5A**) (*39, 43–48*). Less than 22% of these genes were also sensitive to NIPBL or WAPL depletion, suggesting that the effect of CTCF knockdown on transcription was mostly distinct from cohesin dysregulation (**fig. S5B**).

However, of the 1,877 genes differentially expressed after NIPBL knockdown, 959 were fully rescued by co-depletion of CTCF (**fig. S5C**). Another 280 genes showed decreased changes in expression; therefore, a total of 66% of DEGs after NIPBL depletion were partially or fully rescued in the double knockdown condition (**fig. S5C**). Interestingly, 85% of the genes rescued by CTCF depletion were also rescued by WAPL depletion, consistent with their shared role in restricting chromatin loop extrusion.

To simultaneously compare all gene expression changes across the six conditions and two biological replicates each, we performed a principal component analysis (PCA) of the PRO-seq datasets (**Fig. 5D**). This analysis reiterates our finding that NIPBL and WAPL depletion had opposing effects on gene expression; these two conditions separated along the second principal component. All replicates for control and NIPBL-WAPL double knockdown conditions were clustered strikingly close to one another, reflecting the genome-wide restoration of transcription observed in these samples. Replicates involving CTCF knockdown were distinctly separated from the other samples along the first principal component, consistent with a large effect on different genes; however, we noted that CTCF samples trended along the second principal component toward samples with depletion of WAPL. Finally, co-depletion with NIPBL did not affect the variance of the first principal component; however, the second principal component reflected the partial rescue of gene expression across all samples. Together, these data strongly support the notion that reduced cohesin activity via NIPBL depletion can be functionally offset by removal of either its negative regulator (WAPL) or the physical barriers (CTCF) that restrict loop-extrusion events.

## Discussion

In this study, we modified levels of the cohesin regulators NIPBL and WAPL to investigate their unique and shared effects on chromatin folding and transcription. Interestingly, ~90% loss of either protein was sufficient to alter chromatin folding by FISH but did not affect mitosis or cell proliferation, indicating that a small amount of NIPBL or WAPL is sufficient for proper sister-chromatin cohesion and chromosome segregation. However, this was not the case for gene regulation considering ~2,000 genes were misexpressed following depletion of either protein.

Given that NIPBL and WAPL are opposing regulators of cohesin (*18, 21, 29, 38*), one prediction might be that each of their knockdowns would alter the same set of genes but in opposite directions. Instead, we found that most (~70%) DEGs were exclusive to either knockdown condition. Moreover, the 30% of DEGs that were shared between the knockdowns tended to be differentially expressed in the same direction. Overall, the DEGs were enriched at cohesin binding sites and anchors of chromatin loops, consistent with their dysregulation due to aberrant looping albeit with differential genomic sensitivities to increased and decreased cohesin activity. Indeed, we found that NIPBL- and WAPL-sensitive genes were both nonrandomly clustered within TADs and coordinately up- or downregulated.

Our results are consistent with a model in which genomic regions are co-regulated within spatial hubs. These hubs could either promote or repress transcription, depending on the local environment (**Fig. 6**) (*16, 49–57*). When NIPBL is depleted, loop extrusion is limited; consequently, distal chromatin may not reach their target regulatory hubs as efficiently, resulting in altered expression of several nearby genes. This is consistent with our analysis of the *MCM5* locus, in which the gene is displaced from neighboring domains following loss of cohesin (*25*). The opposite would occur in the absence of WAPL, with regions beyond those normally incorporated into hubs brought into proximity, providing an explanation for its role in expression of a different set of genes. Therefore, while not essential for gene expression, NIPBL and WAPL may function to balance exposure of promoters within a TAD to local gradients of eRNAs and activated TFs (*58*).

**Fig. 6:**
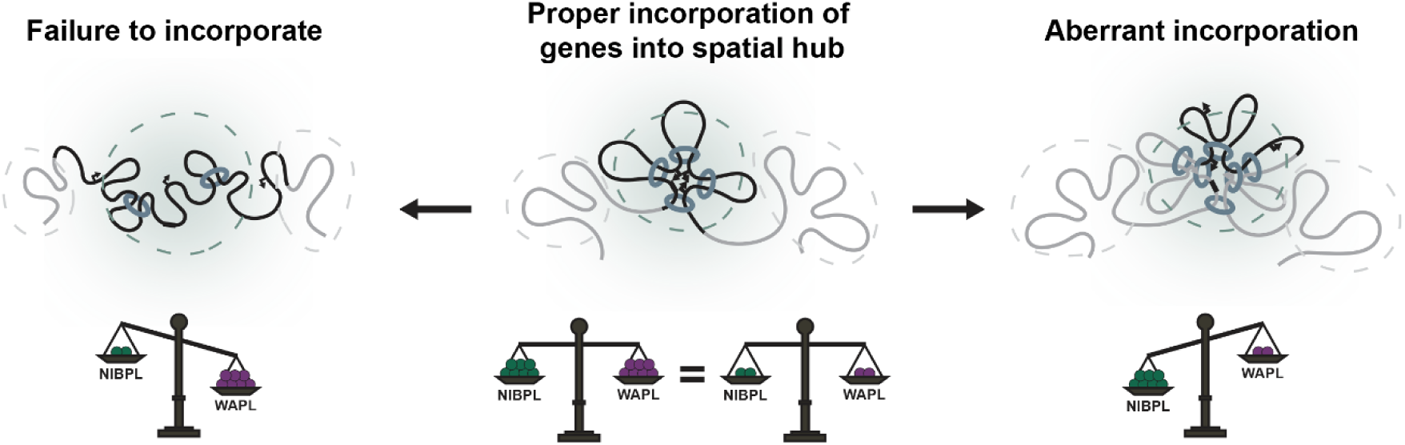
NIPBL and WAPL balance cohesin activity to regulate gene expression. Cohesin normally promotes the compaction and clustering of chromatin into TADs, which we propose act as regulatory hubs. These hubs can either stimulate or repress expression of nearby genes in a dynamic fashion. Depletion of NIPBL would limit loop extrusion events and processivity, resulting in less frequent incorporation of distal genes into their target hubs. In contrast, WAPL depletion would effectively increase loop extrusion events and lead to the ectopic incorporation of distal genes into new hub environments. Co-depletion of NIPBL and WAPL would therefore balance cohesin activity, restore chromatin folding, and correct hub formation to effectively rescue gene expression.

Interestingly, balancing the expression of these two ubiquitously expressed and essential proteins rescued the effects of knockdown of either single protein. In total, ~85% of genes differentially expressed after NIPBL or WAPL knockdown were at least partially rescued by simultaneous knockdown of both proteins to ~10% control levels. Co-depletion also partially restored the levels of chromatin-bound cohesin and rescued the spatial insulation between TADs by FISH. Contact between a boundary-proximal gene sensitive to cohesin loss, *MCM5*, and its neighboring regulatory domain, was also rescued in the double knockdown condition. This was accompanied by correction of *MCM5* expression, which is consistent with its dependency on proper cohesin activity. Indeed, we found that co-depletion of NIPBL and CTCF largely rescued the same DEGs as WAPL. This suggests that proper gene expression is achieved by balancing different restrictions to cohesin-mediated loop extrusion.

Together, our data are in full agreement with several intriguing findings in which co-depletion of WAPL and NIPBL or MAU2 functionally restore proper organismal development, cellular differentiation rates, or cell viability across *Drosophila*, mouse, and human systems (*29, 59, 60*). Here, we show that this rescue extends to the molecular level resulting in near complete complementation of gene expression changes across the entire genome. We therefore propose that the correct stoichiometric ratio, rather than the absolute amount, of NIPBL and WAPL is necessary to properly modulate cohesin activity, organize chromatin, and regulate transcription.

## Materials and Methods

### Experimental design

This study used FISH and PRO-seq to study the consequences of knocking down the cohesin regulatory factors NIPBL and WAPL in the human HCT-116 cell line.

### Cell culture

HCT-116 cells were obtained from AATC (ATCC CCL-247 Colon Carcinoma; Human; Lot 70009735) and HCT-116-RAD21-AID cells were obtained from Natsume et al. (*61*). Cells were cultured in McCoy’s 5A media supplemented with 10% FBS, 2 mM L-glutamine, 100 U/ml penicillin, and 100ug/ml streptomycin and filtered using a 0.22-μm PES membrane at 37°C with 5% CO_2_. HCT-116-RAD21-AID cells were re-selected with 100µg/ml G418 and 100µg/ml HygroGold prior to experiments. Prior to FISH on slides, HCT-116-RAD21-AID cells were synchronized as previously described in (*25*).

### RNAi

The following siRNAs (Dharmacon) were used: Non-targeting control (D-001210-05-05), NIPBL (J-012980-08; target sequence: 5’-CAACAGAUCACAUAGAGUU-3’), WAPL (L-026287-01-0005; target sequences administered as a pool: 5’-GGAGUAUAGUGCUCGGAAU-3’, 5’-GAGAGAUGUUUACGAGUUU-3’, 5’-CAAACAGUGAAUCGAGUAA-3’, 5’-CCAAAGAUACACGGGAUUA-3’), and CTCF (L-020165-00-0010; target sequences administered as a pool: 5’-GAUGAAGACUGAAGUAAUG-3’, 5’-GGAGAAACGAAGAAGAGUA-3’, 5’-GAAGAUGCCUGCCACUUAC-3’, 5’-GAACAGCCCAUAAACAUAG-3’). siRNAs were incubated for 20 minutes at room temperature with RNAiMAX Lipofectamine transfection reagent (ThermoFisher) in Opti-MEM reduced serum media (ThermoFisher) and seeded into wells. HCT-116 cells were trypsinized and resuspended in antibiotic free media (McCoy’s 5A media supplemented with 10% FBS and 2 mM L-glutamine), then plated onto siRNA for a final siRNA concentration of 50 nM (non-targetting control and WAPL), 100 nM (NIPBL), or 150 nM (CTCF). For CTCF knockdowns, cells were retreated with 150nM CTCF siRNAs 24 hours after initial treatment. After 72h (NIPBL, WAPL, non-targeting control) or 96h from the initial RNAi treatment (CTCF), cells were harvested for experiments.

### Western blotting

To prepare samples, cells were trypsinized and resuspended in fresh media, washed once in cold Dulbecco’s PBS, and then centrifuged at 500g for 5 min at 4°C. Subcellular protein fractionations were performed using the Subcellular Protein Fractionation Kit for Cultured Cells (Thermo Scientific, Catalog no: 78840) according to the product manual. We used reagent volumes corresponding to 10μl packed cell volume for 4×10^6^ cells. In step 10, we incubated samples at room temperature for 15 minutes. To extract the whole cell lysate (WCL), samples were resuspended in 1x RIPA buffer with fresh protease inhibitors (200μl per 5×10^6^ cells), nutated 30 min at 4°C, centrifuged at 16,000g for 20 min at 4°C. Supernatant was extracted and stored at −80°C. The Pierce BCA protein assay kit (Catalog no. 23225) was used to quantify the amount of protein per sample.

For each sample, 12-15μg protein was combined with NuPAGE LDS sample buffer and sample reducing agent (Thermo Fisher Scientific). Samples were denatured at 70°C for 10 min, then cooled on ice. Benzonase was added to the WCL samples (0.5μl), followed by a 15-min incubation at 37°C. We 25μl of each sample on Mini-PROTEAN TGX precast gels (Bio-Rad, catalog no. 456-1083) at 35mA. Protein was then transferred to 0.2 µm nitrocellulose membrane at 110 V for 1hr. The nitrocellulose membrane was then washed twice in TBS (150 nM NaCl, 20 mM Tris) for 5 min, and blocked in 5% milk in TBS-T (TBS with 0.05% Tween 20) for 30 min.

The membrane was washed again twice in TBS-T, then incubated with primary antibody diluted in 5% milk in TBS-T overnight at 4°C. The following day, the nitrocellulose membrane was washed twice in TBS-T for 5 min each wash, then incubated with secondary antibodies diluted in 5% milk in TBS-T for 1 h at RT. The nitrocellulose filter was then washed twice in TBS-T for 15 min each wash, followed by a final 15-min wash in TBS. For blots probed with secondary antibodies conjugated to horseradish peroxidase (HRP), the membrane was incubated in a 1:1 mixture of Clarity Western ECL Substrate reagents (Bio-Rad). Blots were then imaged on a ChemiDoc MP Imaging System and analyzed with Bio-Rad Image Lab software (v6.1.0 build 7). The following primary antibodies were used: NIPBL (sc-374625; 1:400), WAPL (Cell Signaling Technology (CST) D9J1U; 1:1,000), RAD21 (ab992, 1:1,000), HDAC2 (Cell Signaling Technology 5113S, 1:2,000), GAPDH (CST 5174S, 1:2,000), H3 (ab1791, 1:4,000). The following secondary antibodies were used: anti-mouse IgG, HRP-linked Antibody (CST #7076; 1,5,000), anti-rabbit IgG, HRP-linked antibody (CST #7074; 1,5,000), Cy3 AffiniPure Goat Anti-Rabbit IgG (Jackson ImmunoResearch 111-165-003, 3:20,000), IRDye 800CW Goat anti-Mouse IgG Secondary Antibody (LI-COR, 3:10,000).

### RNA extraction and RT-qPCR

HCT-116 derived RNA was isolated using the RNeasy Plus Kit (Qiagen) according to manufacturer’s instructions. For complementary DNA (cDNA) synthesis, a 50μl reaction containing 20μl RNA, 1500pmol Oligo dT primer (IDT), 1.6mM dNTPs, 1x RT Buffer (Thermo Scientific), 0.5μl RNase OUT (Invitrogen), and 0.7μl Maxima RT (Thermo Scientific) was incubated at 50°C for two hours then 85°C for 5 min. Samples were stored at −20°C until use. RT-PCR was performed using PowerUP Sybr (ThermoFisher, #A25741) based on manufacturer’s instructions. Briefly, cDNA was diluted to a working concentration of 6μg and HCT-116 genomic DNA (gDNA) was diluted in a 1:10 serial dilution. A 6μl reaction was prepared per well, with 1x PowerUP Sybr and 0.2μM of the forward and reverse primers and combined with 4μl diluted DNA. Each reaction was performed in triplicate. qPCR was performed on the QuantStudio7 Flex System. YWHAZ and TBP were used as reference control genes. The sequences of oligonucleotides used for qPCR are: NIPBL forward primer: 5’-TCTCTTTGTTACTTGTCTGTTTCC-3’ and reverse primer 5’-ATGTTTTGCTTTGAAAACCAGTG-3’; WAPL forward primer 5’-GAACTAAAACAGCTCCATCACC-3’ and reverse primer 5’-CACACTTTCAGGCACACCAG-3’; YWHAZ forward primer 5’-CCCGTTTCCGAGCCATAAAAG-3’ and reverse primer 5’-TTTGGCCTTCTGAACCAGCTC-3’; and TBP forward primer 5’-ACAGCTCTTCCACTCACAGAC-3’ and reverse primer 5’-ATGGGGGAGGGATACAGTGG-3’.

### FISH Probe design & synthesis

Oligopaint probes were designed as previously described (*25*). Briefly, we designed probes to domains and subdomains based on ChIP-Seq and Hi-C data using the OligoMiner design pipeline (*62*). Probe coordinates and details are listed in Table 8. Oligopaints were designed to have either 80 bases of homology and were purchased from Twist Bioscience. Additional bridge probes were designed to the *MCM5* gene probe to amplify its signal (*63*). Oligopaints were synthesized as previously described (*25*) with some modifications to allow for direct conjugation to fluorescent dyes. Specifically, aminoallyl-dUTP (ThermoFisher Scientific) was incorporated into the probes to allow for conjugation with Alexa 488 (ThermoFisher Scientific), Cy3 (Gold Biotechnology), or Alexa 647 (ThermoFisher Scientific).

### DNA Fluorescence in situ hybridization (FISH)

#### FISH on Slides

FISH was performed on slides to chr2: 217-222Mb (**Fig. 1, J to L and fig. S1, G to I**) and chr22: 33-36.2Mb (**Fig. 4, E to G and I to K**). Cells were settled on poly(L-lysine)-treated glass slides for 2 h. Cells were then fixed to the slide or coverslip for 10 min with 4% formaldehyde in phosphate-buffered saline (PBS) with 0.1% Tween 20, followed by three washes in PBS for 5 min each wash. Slides and coverslips were stored in PBS at 4°C until use. Prior to FISH, slides were warmed to room temperature (RT) in PBS for 10 min. Cells were permeabilized in 0.5% Triton-PBS for 15 min. Cells were then dehydrated in an ethanol row, consisting of 2-min consecutive incubations in 70%, 90% and 100% ethanol. The slides were then allowed to dry for about 2 min at RT. Slides were incubated for 5 min each in 2xSSCT (0.3 M NaCl, 0.03 M sodium citrate and 0.1% Tween 20) and 2xSSCT/50% formamide at RT, followed by a 1-h incubation in 2xSSCT/50% formamide at 37°C. Hybridization buffer containing primary Oligopaint probes, hybridization buffer (10% dextran sulfate, 2xSSCT, 50% formamide and 4% polyvinylsulfonic acid (PVSA)), 5.6 mM dNTPs and 10 µg RNase A was added to slides, covered with a coverslip, and sealed with rubber cement. 50 pmol of probe was used per 25 µl hybridization buffer. Slides were then denatured on a heat block in a water bath set to 80°C for 30 min, then transferred to a humidified chamber and incubated overnight at 37°C. The following day, the coverslips were removed and slides were washed in 2xSSCT at 60°C for 15 min, 2xSSCT at RT for 10 min, and 0.2xSSC at RT for 10 min. Next, hybridization buffer (10% dextran sulfate, 2xSSCT, 10% formamide and 4% PVSA) containing secondary probes conjugated to fluorophores (10pmol per 25 µl buffer) was added to slides, covered with a coverslip and sealed with rubber cement. Slides were placed in a humidified chamber and incubated for 2 h at RT. Slides were washed in 2xSSCT at 60°C for 15 min, 2xSSCT at RT for 10 min, and 0.2xSSC at RT for 10 min. To stain DNA, slides were washed with Hoechst (1:10,000 in 2xSSC) for 5 min. Slides were then mounted in SlowFade Gold Antifade (Invitrogen).

Images were acquired on a Leica widefield fluorescence microscope, using a 1.4 NA ×63 oil-immersion objective (Leica) and Andor iXonµltra emCCD camera. All images were deconvolved with Huygens Essential v20.04.03 (Scientific Volume Imaging), using the CMLE algorithm, with a signal to noise ratio of either 40, and 40 iterations (DNA FISH) or signal to noise ratio of 40 and 2 iterations (DNA stain). The deconvolved images were segmented and measured using a modified version of the TANGO 3D-segmentation plug-in for ImageJ (*64–66*). Edges of nuclei and FISH signals were segmented using a Hysteresis-based algorithm.

#### High-throughput DNA or RNA Oligopaints (HiDRO)

All other FISH experiments (**Fig. 1M, fig. S1, L and M**, **Fig. 4H**, and **fig. S4G)** were performed using HiDRO as described in detail in (*41*). All spins were performed at 1200 rpm for 2 min at RT unless otherwise indicated. When possible, automatic pipetting was performed by a Matrix WellMate (Thermo Fisher Scientific). For experiments in the HCT-116-RAD21-AID cell line, 7.5×10^4^ cells in supplemented McCoy’s 5A media −/+ 500 µM auxin were seeded in 384-well plates (Perkin Elmer 6057300) and incubated at 37°C for 6 h. For RNAi experiments in the HCT-116 cell line, plates were seeded with siRNA (see RNAi section for details) diluted in Opti-MEM reduced serum medium to a final concentration of 25nM per well. Plates were then spun and incubated at RT for 20 min. HCT-116 cells were trypsinized and resuspended in antibiotic-free medium, then 2.5×10^3^ cells were seeded in each well. Plates were spun and incubated at 37°C for 72 h.

Following incubation, media was aspirated, all wells had PBS added to them, and plates were spun. PBS was aspirated and cells were fixed in each well with 4% PFA, 0.1% Tween-20 in 1x PBS for 10 minutes at RT. Plates were spun once during fixation. Then plates were rinsed with 1xPBS and washed twice for 5 minutes with 1xPBS with a spin during each wash. 70% ethanol was then added to each well, plates were sealed with foil plate covers (Corning) and stored for at least 20 hours at 4°C until used for FISH.

On the first day of DNA FISH, ethanol was aspirated and plates were washed in 1xPBS for 10 min to reach RT. Plates were then spun, washed briefly again in 1xPBS and spun again. Cells were permeabilized for 15 min in 0.5% Triton-X and 5 minutes in 2xSSCT (0.3 M NaCl, 0.03 M sodium citrate and 0.1% Tween 20). Then 2xSSCT/50% formamide was added to all wells, and plates were double sealed with foil covers. Pre-denaturation was performed at 91°C for 3 min and then 60°C for 20 min on heat blocks (VWR). After plates were spun, foil covers were removed and hybridization mix was added to wells. Hybridization mix consisted of 50% formamide, 10% dextran sulfate, 4% PVSA, 0.1% Tween-20, 2xSSC, and each probe at 0.1pmol/µl. 2pmol of probe was used per 20µl of hybridization mix. Hybridization mix was viscous and thus pipetted using a manual multichannel pipette. After spinning, plates were double sealed with foil covers and denatured at 91°C for 20 min on heat blocks. Heat blocks were covered to block light and preserve primary fluorescently labeled probes. Plates were spun after denaturation and then hybridized overnight at 37°C.

The following day, hybridization mix was aspirated, and plates were washed quickly twice with RT 2x SSCT, then with 60°C 2xSSCT for 5 min. Plates were then washed with RT 2x SSCT for 5 min. Nuclei were stained by washing for 5 min in Hoescht (1:10,000 in 2x SSCT). Plates were spun, washed for 15 min with 2x SSC and spun again. Finally, plates were mounted with imaging buffer (2x SSC, 10% glucose, 10mM Tris-HCl, 0.1 mg/ml catalase, 0.37 mg/ml glucose oxidase) and imaged within 5 days of FISH.

Images for HiDRO experiments were acquired on a Molecular Devices Image Xpress Micro 4 Confocal high-content microscope with 0.42 um pinhole and 1.4 NA 60X water immersion objective (Molecular Devices). Max projections of z-series (6 images, 0.5 uM spacing) were generated automatically in MetaXpress and used for downstream analyses.

### Hi-C analysis

Loops were called using the HICCUPS tool in Juicer (version 1.22.01) using the same settings as (*67*) for high resolution maps, as shown here: “-k KR -f .1,.1 -p 4,2 -i 7,5 -t 0.02,1.5,1.75,2 -d 20000,20000”.

TAD were called using the hicFindTAD function of the HiCExplorer package (version 3.7.2) (*68–70*). First, .hic files were first converted to .cool files at 50 kb resolution using hic2cool (https://github.com/4dn-dcic/hic2cool) and then corrected used the “cooler balance” function from the cooltools package (https://github.com/open2c/cooler) (*71*) (Abdennur and Mirny, 2020). These .cool files were then converted to .h5 format using “hicConvertFormat” from HiCExplorer package, and the resulting ..h5 files were used to call TADs with the following parameters of hicFindTADs: “--correctForMultipleTesting fdr --minBoundaryDistance 100000 --delta 0.4”.

### Permutation analyses

Permutation analysis was used to create an “expected” null distribution with which to compare the observed clustering and coordination of DEGs. Most (95%) active genes were within a called TAD. For clustering, all genes in the genome were either assigned transcription status (active/non-active) or the DEG status (DEG/nonDEG). Observed clustering was calculated by measuring the percentage of active genes/DEGs per TAD and comparing it against a 1000 random permutations, where the transcription/DEG status was shuffled across all genes for each permutation while keeping the number of genes in each category constant. A p-value was reported as the percentile ranking of the observed clustering against this permutation distribution. For analysis of the coordination of DEGs within TADs, the same approach was taken as above, with each DEG assigned a direction of misexpression (up/down) and the observed coordination across TADs compared against 1000 random permutations.

### PRO-seq & analysis

#### Cell permeabilization

RNAi was performed as previously described. Following 72 h knockdown, cells were rinsed with Dulbecco’s phosphate buffered saline (DPBS) and treated with trypsin to detach them from the plate. Cells were resuspended in cold supplemented McCoy’s 5A media and three wells of a 6-well plate were pooled per replicate and placed on ice. From this point on, all steps were performed on ice, all buffers were pre-chilled, and samples were spun at 300xg for 10 min at 4°C, unless otherwise noted. Cells were rinsed in PBS containing 1% Bovine Serum Albumin (BSA) to prevent cell clumping, and then resuspended in 1 ml Buffer W (10 mM Tris-HCl pH 8.0, 10 mM KCl, 250 mM sucrose, 5 mM MgCl2, 0.5 mM DTT, 10% glycerol, 1% BSA) then strained through a 35 μm nylon mesh filter. The tube was rinsed with an additional 1ml of Buffer W and passed through the same strainer. A 9X volume of Buffer P (Buffer W + 0.1% IGEPAL CA-630) was immediately added to each sample and nutated for 2 minutes at room temperature. Cells were resuspended in 500μl Buffer F (50 mM Tris-Cl pH 8.3, 40% glycerol, 5 mM MgCl2, 0.5 mM DTT, 1 μL/ml SUPERaseIn RNase inhibitor, 0.5% BSA) using a wide-bore P1000 tip and transferred to a low binding tube. The original tube was rinsed with another 500μl Buffer F and the samples were pooled. Samples were spun at 400xg and resuspended to 5 x 10^6^ cells in 500μl Buffer F. Samples were flash frozen in liquid nitrogen and stored at −80°C

#### PRO-seq library construction

PRO-seq library construction and data analysis was performed by the Nascent Transcriptomics Core at Harvard Medical School, Boston, MA. Aliquots of frozen (−80°C) permeabilized cells were thawed on ice and pipetted gently to fully resuspend. Aliquots were removed and permeabilized cells were counted using a Luna II, Logos Biosystems instrument. For each sample, 1 million permeabilized cells were used for nuclear run-on, with 50,000 permeabilized *Drosophila* S2 cells added to each sample for normalization. Nuclear run on assays and library preparation were performed essentially as described in Reimer et al. (*72*) with modifications noted: 2X nuclear run-on buffer consisted of (10 mM Tris (pH 8), 10 mM MgCl2, 1 mM DTT, 300mM KCl, 40uM/ea biotin-11-NTPs (Perkin Elmer), 0.8U/μl SuperaseIN (Thermo), 1% sarkosyl). Run-on reactions were performed at 37°C. Adenylated 3’ adapter was prepared using the 5’ DNA adenylation kit (NEB) and ligated using T4 RNA ligase 2, truncated KQ (NEB, per manufacturers instructions with 15% PEG-8000 final) and incubated at 16°C overnight. 180μl of betaine blocking buffer (1.42g of betaine brought to 10ml with binding buffer supplemented to 0.6 uM blocking oligo (TCCGACGATCCCACGTTCCCGTGG/3InvdT/)) was mixed with ligations and incubated 5 min at 65°C and 2 min on ice prior to addition of streptavidin beads. After T4 polynucleotide kinase (NEB) treatment, beads were washed once each with high salt, low salt, and blocking oligo wash (0.25X T4 RNA ligase buffer (NEB), 0.3uM blocking oligo) solutions and resuspended in 5’ adapter mix (10 pmol 5’ adapter, 30 pmol blocking oligo, water). 5’ adapter ligation was per Reimer but with 15% PEG-8000 final. Eluted cDNA was amplified 5-cycles (NEBNextµltra II Q5 master mix (NEB) with Illumina TruSeq PCR primers RP-1 and RPI-X) following the manufacturer’s suggested cycling protocol for library construction. A portion of preCR was serially diluted and for test amplification to determine optimal amplification of final libraries. Pooled libraries were sequenced using the Illumina NovaSeq platform.

#### PRO-seq data analysis

All custom scripts described herein are available on the AdelmanLab Github (https://github.com/AdelmanLab/NIH_scripts). Using a custom script (trim_and_filter_PE.pl), FASTQ read pairs were trimmed to 41bp per mate, and read pairs with a minimum average base quality score of 20 retained. Read pairs were further trimmed using cutadapt 1.14 to remove adapter sequences and low-quality 3’ bases (--match-read-wildcards -m 20 -q 10). R1 reads, corresponding to RNA 3’ ends, were then aligned to the spiked in Drosophila genome index (dm3) using Bowtie 1.2.2 (-v 2 -p 6 --best --un), with those reads not mapping to the spike genome serving as input to the primary genome alignment step (using Bowtie 1.2.2 options -v 2 --best). Reads mapping to the hg38 reference genome were then sorted, via samtools 1.3.1 (-n), and subsequently converted to bedGraph format using a custom script (bowtie2stdBedGraph.pl). Because R1 in PRO-seq reveals the position of the RNA 3’ end, the “+” and “-“ strands were swapped to generate bedGraphs representing 3’ end position at single nucleotide resolution.

For a table of statistics, including raw read counts, mappable read counts to the spike in and reference genomes, refer to Table S9. Pairwise correlation (Spearman’s rho) of counts in windows ±2kb around filtered TSS annotation noted in Table S10.

For promoter reads, annotated transcription start sites were obtained from Ensembl v99 for hg38. After removing transcripts with {immunoglobulin, Mt_tRNA, Mt_rRNA} biotypes, PRO-seq signal in each sample was calculated in the window from the annotated TSS to +150 nt downstream, using a custom script, make_heatmap.pl.

Given good agreement between replicates (Spearman’s rho ≥0.95) and similar return of spike-in reads, bedGraphs were merged within conditions, and depth-normalized, to generate bigWig files binned at 10bp.

To determine differentially expressed genes in PRO-seq analyses, the 5’ ends from all PRO-seq reads were used to identify active transcription start sites using a custom script, proTSScall available on the NascentTranscriptionCore GitHub (https://github.com/NascentTranscriptionCore/proTSScall). Briefly, PRO-seq 3’ read bedGraphs for “+” and “-“ strands were separately combined across samples and the composite read counts were assigned to TSS-proximal windows (TSS to +150nt) using the same filtered TSS annotation described above. TSSs with ≤9 counts in this window are deemed ‘inactive’ and the remaining TSSs, deemed ‘active’, are collapsed to yield 1 dominant TSS per gene, defined as the one with the highest TSS-proximal read count -- if the highest read count is shared amongst multiple transcripts, the TSS furthest upstream, in a strand-aware fashion, is called dominant. Dominant TSSs sharing the same start position are deduplicated as follows: (1) if start positions are equal, the TSS with the longest associated annotated transcript is called dominant, (2) if start positions and transcript lengths are both equal, the TSS associated with the lowest Ensembl gene ID (numerical portion) is dominant.

#### Principle component analysis

PRO-seq 3’ reads were summed across the 2kb downstream of each TSS and genes with non-zero sums in at least one sample were retained for PCA analysis. The PCA was generated with the plotPCA function within DESeq2 using the rlog-transformed sums.

#### dREG Enhancer Peak Calling

Enhancer peaks were called using the dREG pipeline (*73*) on merged PRO-seq bigwigs using the default parameters. Peaks were filtered by p-value of 0.02 or less and dREG score of 0.55 or more. Resulting peaks list was manually curated into standard bed format. Centers called outside of the dREG peak area were manually moved to the closest end of the dREG peak. dREG scores were multiplied by 1000 and converted to integers to conform to standard BED file format. Peaks assigned an “NA” p-value from DESeq2 were removed (v1.30.1) (*74*). Promoter proximal dREG peaks within 1kb of an annotated TSS (Ensembl v99) were filtered using the UCSC Table Browser (*75*). All other peaks were annotated as “distal”. Intragenic peaks were defined as distal dREG peaks that overlapped an annotated gene body. All others were flagged as intergenic.

#### Differential Expression Analysis

Differential expression analysis was performed in R v3.6.1 with DESeq2 v1.30.1 (*74*). Read counts were obtained over whole genes from TSS to TES as defined by proTSScall, distal dREG peaks, TSS proximal regions (dominant TSS to TSS+150bp), and gene bodies (dominant TSS+250 to TSS+2250bp) using the featureCounts function from Rsubread v1.34.7 (*76*). Defaults were used with the following exceptions: minMQS=10; countChimericFragments=FALSE; isPairedEnd=FALSE; strandSpecific=2 (or strandSpecific=0 for distal dREG peaks); nthreads=8. DESeq2 was run with defaults using the nbinomWaldTest function. The size factors obtained from whole gene bodies were applied to all other groups. Log fold change shrinkage was performed using the ‘apeglm’ algorithm (*77*). Significant differentially expressed genes were filtered for a minimum adjusted p-value of 0.01 or less, removing NA values.

## Statistical Analysis

The numbers of samples (n), p values, and specific statistical tests performed for each experiment are noted in the figure legends. Biological replicates involved an independent isolation of cells including any relevant treatment. HiDRO replicates represent separate wells of a 384-well plate. Statistical analyses were performed using Prism 9 software by GraphPad (v9.2.0).

## Acknowledgments

We would like to thank Leah Rosin and members of the Joyce laboratory for helpful discussions and critical reading of the manuscript. In addition, we thank M. Kanemaki for the HCT-116-RAD21-AID cell line and the Nascent Transcriptomics Core at Harvard Medical School, Boston, MA for performing PRO-seq library construction and for assistance with data analysis.

## Author contributions

JML and EFJ designed the study. JML, AF, SCN, DSP, YL, and EFJ analyzed the results. JML, DSP, and PS performed the experiments. SCN and RY generated the Oligopaint probes used in this study. JML and EFJ wrote the manuscript. All authors discussed the results and commented on the manuscript.

## Data and materials availability

All data needed to evaluate the conclusions in the paper are present in the paper and/or the Supplementary Materials or are available from the corresponding author upon request. Raw and processed PRO-seq (GSE200773), Hi-C and ChIP-seq data (GSE199607) were deposited into the GEO database. Hi-C data were re-analyzed as described in the methods.

## Supplemental Figures

**Figure S1:**
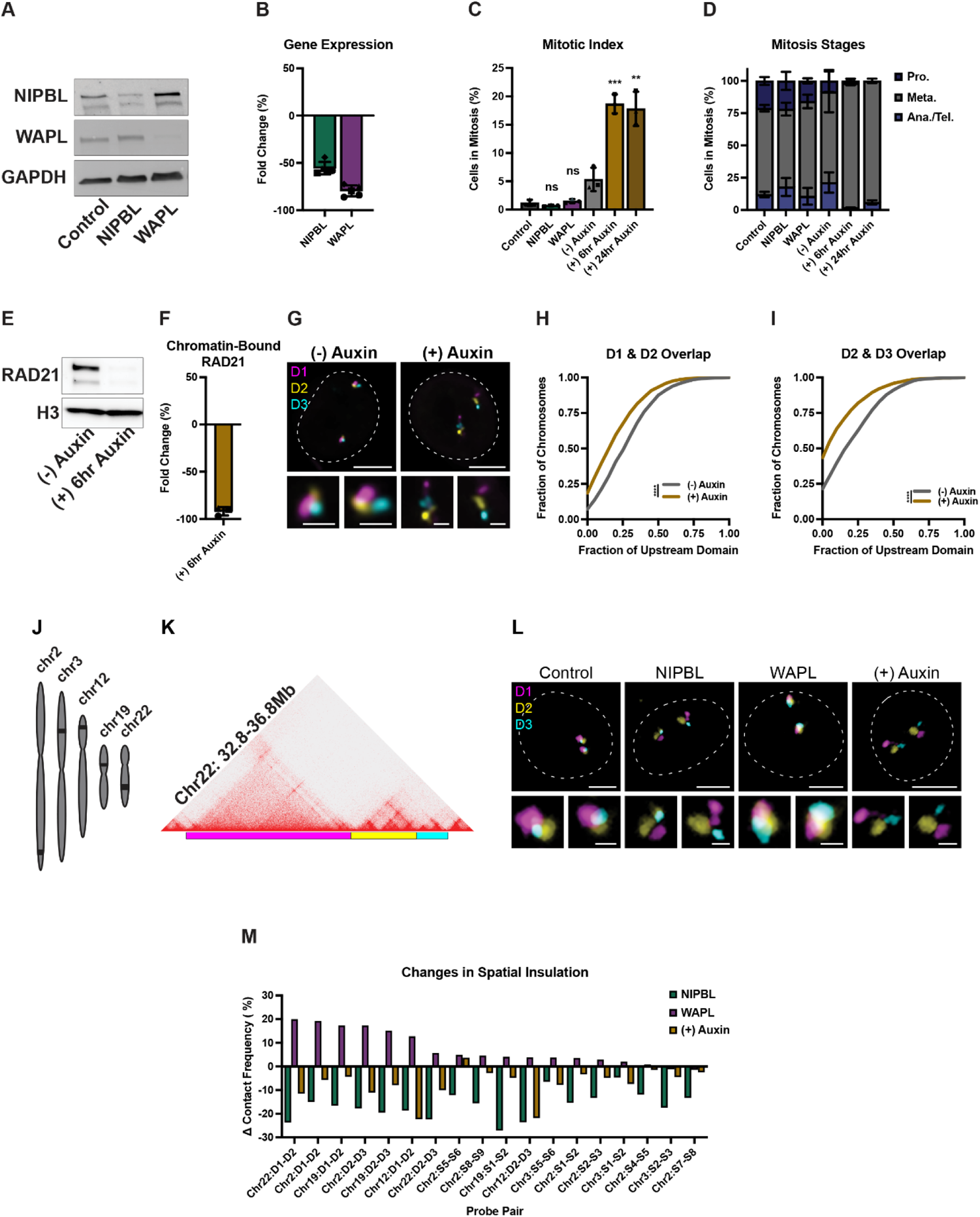
Additional information related to Figure 1. (**A**) Fluorescent western blot to NIPBL (top of the two bands) and WAPL in the whole cell lysate from RNAi control, NIPBL, or WAPL depleted HCT-116 cells. (**B**) Mean fold change (%) in expression by qPCR for NIPBL and WAPL in each respective knockdown. Each symbol represents a biological replicate, error bars represent standard deviation. (**C**) Mitotic index measured by percentage of cells that stained positive for phospho-Histone H3 (PH3) by IF in RNAi control, NIPBL, or WAPL depleted HCT-116 cells and HCT-116-RAD21-AID cells −/+ auxin for 6 or 24 hours. Each bar represents the mean of 3 biological replicates, error bars represent standard deviation. Unpaired t test, *** p < 0.001, ** p = 0.004, ns = not significant (p = 0.23 for Control vs. NIPBL; p = 0.44 for Control vs. WAPL). (**D**) Average percentage of mitotic cells in each stage of mitosis in RNAi control, NIPBL, or WAPL depleted HCT-116 cells and HCT-116-RAD21-AID cells −/+ auxin for 6 or 24 hours. Pro. = prometaphase, Meta. = metaphase, Ana./Telo. = Anaphase or Telophase. Each bar represents the average of 3 biological replicates, error bars represent standard deviation. (**E**) HRP western blot to RAD21 in chromatin-bound subcellular protein fractionations of HCT-116-RAD21-AID cells −/+ auxin for 6 hours. All bands from the same blot. (**F**) Mean fold change (%) of RAD21 bound to chromatin in HCT-116-RAD21-AID cells −/+ auxin. Each symbol represents a biological replicate, error bars represent standard deviation. (**G**) Representative FISH images for three domains at chr2:217-222Mb in HCT-116-RAD21-AID cells −/+ auxin. Dashed line represents nuclear edge, scale bar, 5μm (above) or 1μm (below). (**H**) Cumulative frequency distribution of overlap between the neighboring domains D1 and D2 on chr2 in HCT-116-RAD21-AID cells before (n = 1,874 chromosomes) and after auxin treatment (n = 2,128 chromosomes). Two-tailed Mann-Whitney test, *** p < 0.001. (**I**) Cumulative frequency distribution of overlap between the neighboring domains D2 and D3 on chr2 in HCT-116-RAD21-AID cells before (n = 1,898 chromosomes) and after auxin treatment (n = 2,190 chromosomes). Two-tailed Mann-Whitney test, *** p < 0.001. (**J**) Chromosome schematic representing the relative locations of the HiDRO Oligopaint FISH probes. (**K**) Oligopaint design for three neighboring domains at chr2:217-222Mb. (**L**) Representative FISH images for three domains at chr2:217-222Mb in HCT-116-RAD21-AID cells −/+ auxin. Dashed line represents nuclear edge, scale bar, 5μm (above) or 1μm (below). (**M**) Change in contact frequency across 18 domain pairs in NIPBL, or WAPL depleted HCT-116 cells and auxin treated HCT-116-RAD21-AID cells. Each bar represents the median of ≥ 4 biological replicates. D indicates domain boundary, S indicates sub-domain boundary.

**Figure S2:**
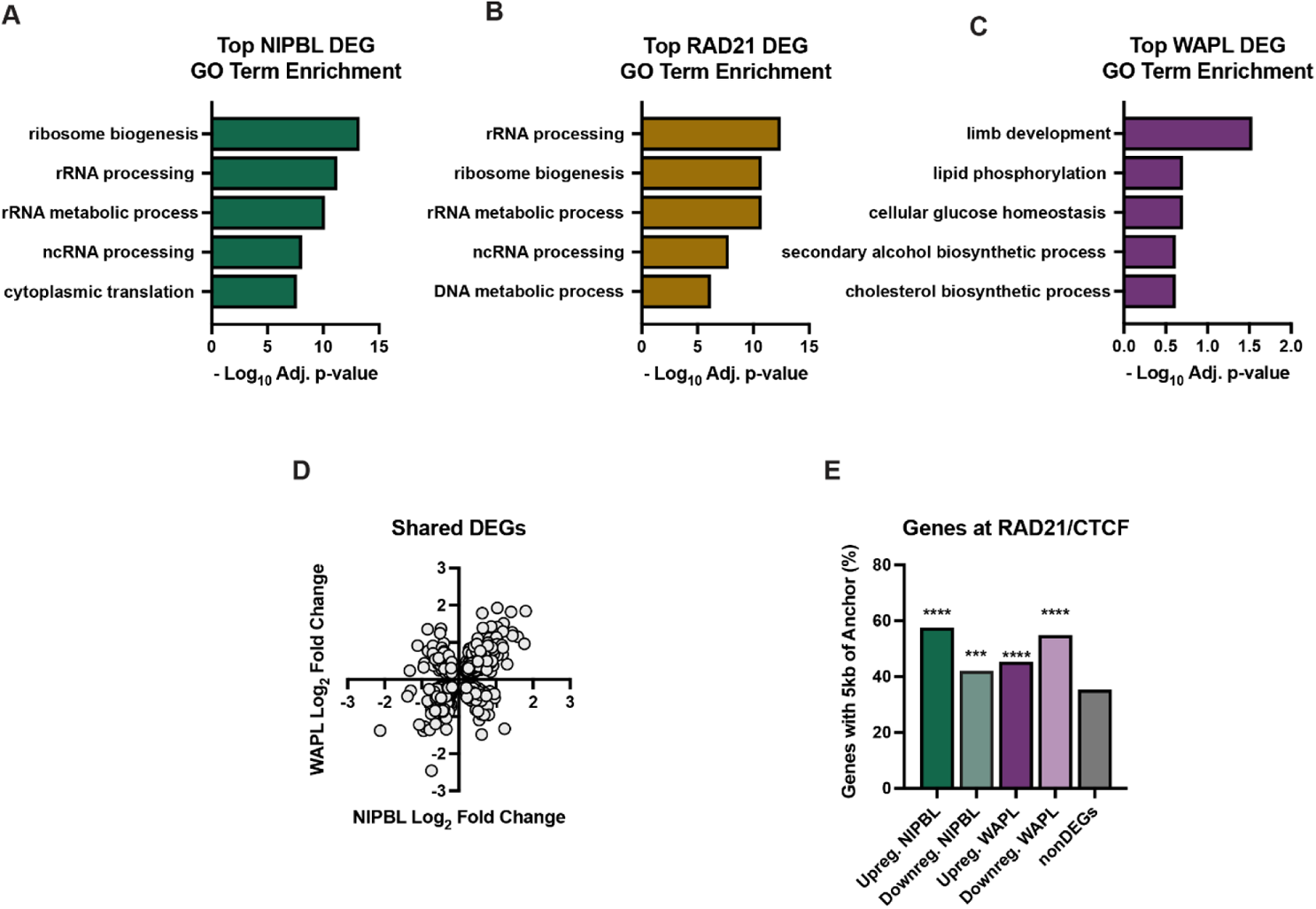
Additional information related to Figure 2. (**A**) Top 5 GO Biological Processes scored by adjusted p-value for NIPBL DEGs and their significance. (**B**) Top 5 GO Biological Processes scored by adjusted p-value for WAPL DEGs and their significance. (**C**) Top 5 GO Biological Processes scored by adjusted p-value for RAD21 DEGs and their significance. (**D**) The log_2_(fold change) of shared DEGs across NIPBL and WAPL knockdown conditions. (**E**) Percentage of up, down, NIPBL, WAPL, or nonDEGs with a TSS within 5kb of a RAD21 ChIP-Seq peak co-occupied by CTCF. Fisher’s exact test, **** p < 0.0001, *** p = 0.0002.

**Figure S3:**
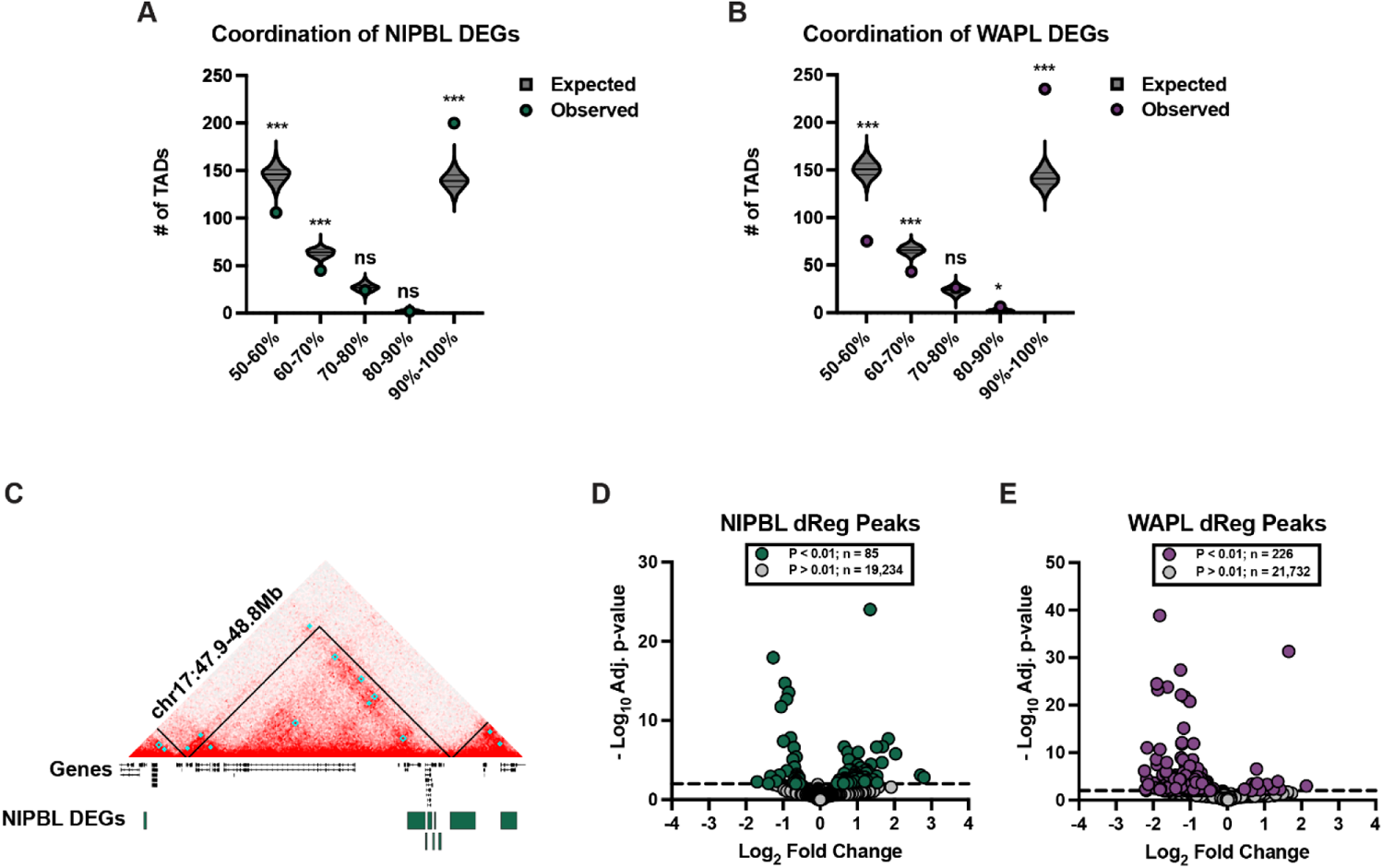
Additional information related to Figure 3. (**A**) The number of expected versus observed TADs with binned coordination scores. 50% coordination represents random misexpression of NIPBL DEGs and 100% coordination represents all NIPBL DEGs in the TAD being up or down regulated. The dot represents the observed data, compared to the expected data in the null distribution (violin plot) generated by shuffling the fold change values amongst the DEGs 1,000 times. (**B**) The number of expected versus observed TADs with binned coordination scores. The dot represents the observed data, compared to the expected data in the null distribution (violin plot) generated by shuffling the fold change values amongst the DEGs 1,000 times. (**C**) Representative TAD with 100% DEG coordination on chr17 which contains nine upregulated NIPBL DEGs. Black lines represent TADs, cyan boxes represent loops. (**D**) The log_2_(fold change) of dREG peaks after NIPBL knockdown versus their significance. Significantly changed dREG peaks are in green (n = 85) and non-significantly changed dREG peaks (adjusted p-value > 0.01) are in grey (n = 19,234). (**E**) The log_2_(fold change) of dREG peaks after WAPL knockdown versus their significance. Significantly changed dREG peaks are in green (n = 226) and non-significantly changed dREG peaks (adjusted p-value > 0.01) are in grey (n = 21,732).

**Figure S4:**
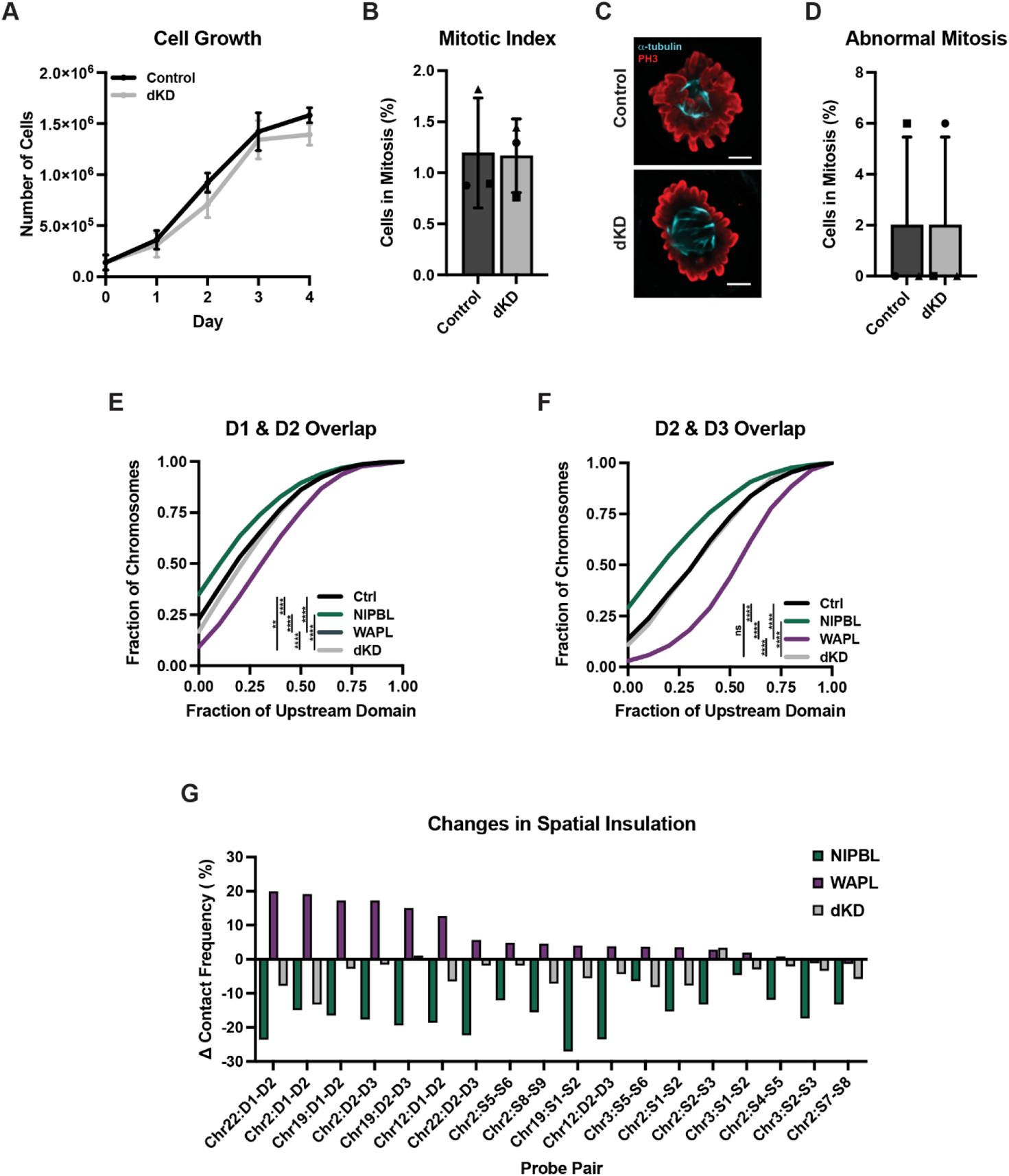
Additional information related to Figure 4. (**A**) Cell growth measured in 24-hour increments following RNAi to NIPBL and WAPL or a non-targeting sequence as the control. Each bar represents the mean of 3 biological replicates and error bars represent the standard deviation. (**B**) Mitotic index measured by percentage of cells that stained positive for phospho-Histone H3 (PH3) by IF in RNAi control or NIPBL and WAPL depleted HCT-116 cells. Each bar represents the mean of 3 biological replicates, error bars represent standard deviation. Unpaired t test, ns = not significant (p = 0.94). (**C**) Representative immunofluorescence images of mitotic cells stained for α-tubulin (cyan) and phospho-Histone H3 (PH3; red) in RNAi control or NIPBL and WAPL depleted HCT-116 cells. Scale bar, 5μm. (**D**) Average percentage of mitotic cells with abnormal mitosis in RNAi control or NIPBL and WAPL depleted HCT-116 cells. Each symbol represents a biological replicate, error bars represent standard deviation. (**E**) Cumulative frequency distribution of overlap between the neighboring domains D1 and D2 on chr2 in RNAi control (n = 1,954 chromosomes), NIPBL (n = 1,584 chromosomes), WAPL (n = 1,677 chromosomes), or dKD (n = 1,711 chromosomes) depleted HCT-116 cells. Two-tailed Mann-Whitney test, **** p < 0.0001, ** p = 0.0012. Biological replicate of data in Fig. 4G. (**F**) Cumulative frequency distribution of overlap between the neighboring domains D2 and D3 on chr2 in RNAi control (n = 1,956 chromosomes), NIPBL (n = 1,671 chromosomes), WAPL (n = 1,666 chromosomes), or dKD (n = 1,728 chromosomes) depleted HCT-116 cells. Two-tailed Mann-Whitney test, **** p < 0.0001, ns = not significant (p = 0.18). Biological replicate of data in Fig. 4I. (**G**) Change in contact frequency across 18 domain pairs in HCT-116 cells depleted for NIPBL, WAPL, or both. Each bar represents the median of ≥ 4 biological replicates. D indicates domain boundary; S indicates sub-domain boundary.

**Figure S5:**
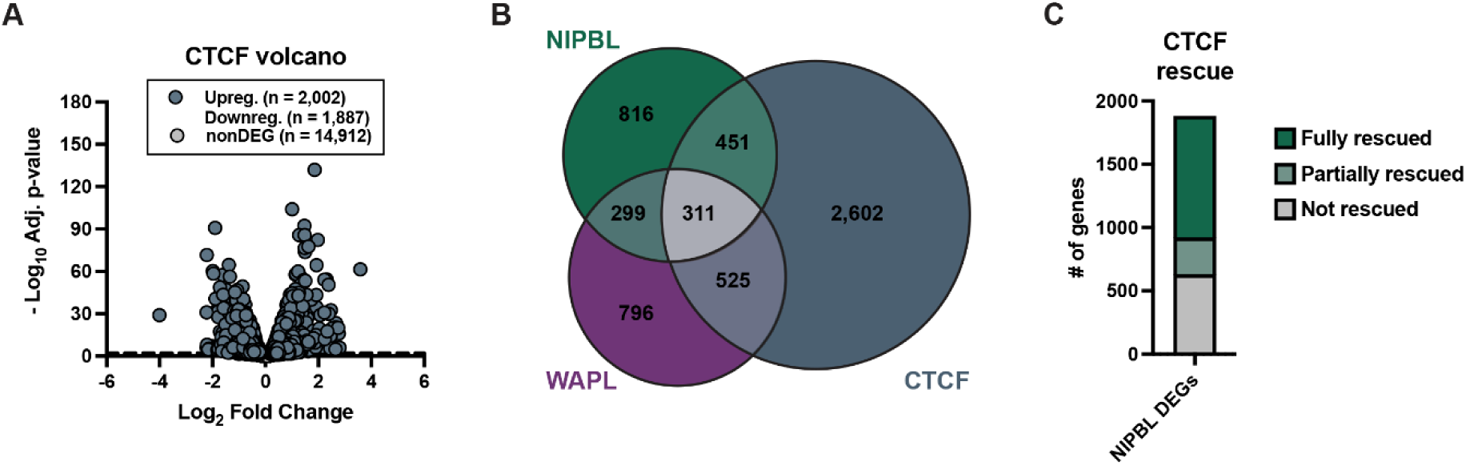
Additional information related to Figure 5. (**A**) The log_2_(fold change) of genes after CTCF knockdown versus their significance. DEGs are in blue (2,002 up, 1,887 down) and non-significantly changed genes (adjusted p-value > 0.01) are in grey. (**B**) Venn diagram of the NIPBL, WAPL, and CTCF DEGs. (**C**) Number of NIPBL DEGs fully, partially, or not rescued in the NIPBL/CTCF double knockdown condition.

## Tables

**Table S1.**
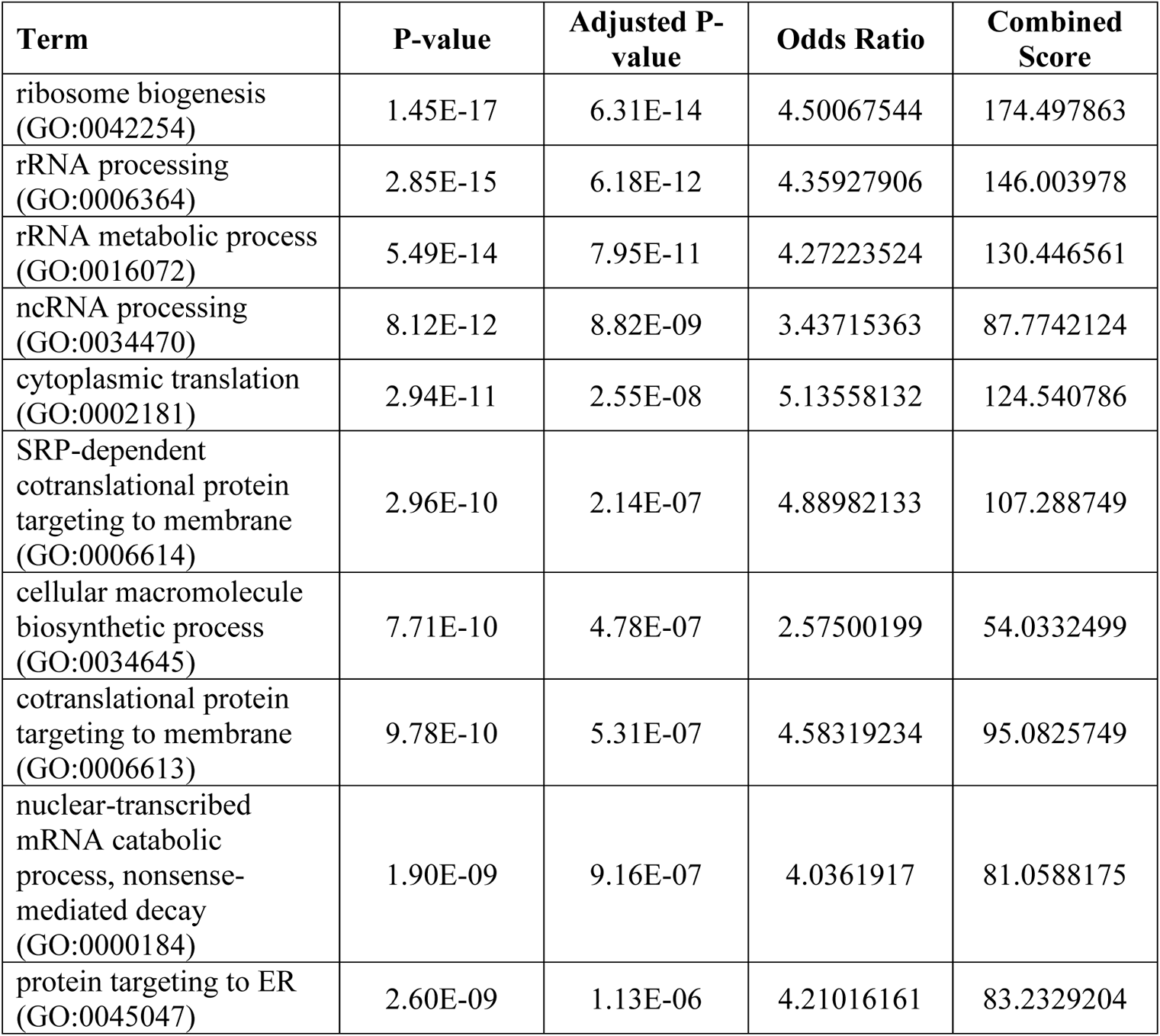
Biological processes associated with NIPBL knockdown. Top 10 GO Biological Processes for NIPBL DEGs sorted by adjusted p-value.

**Table S2.**
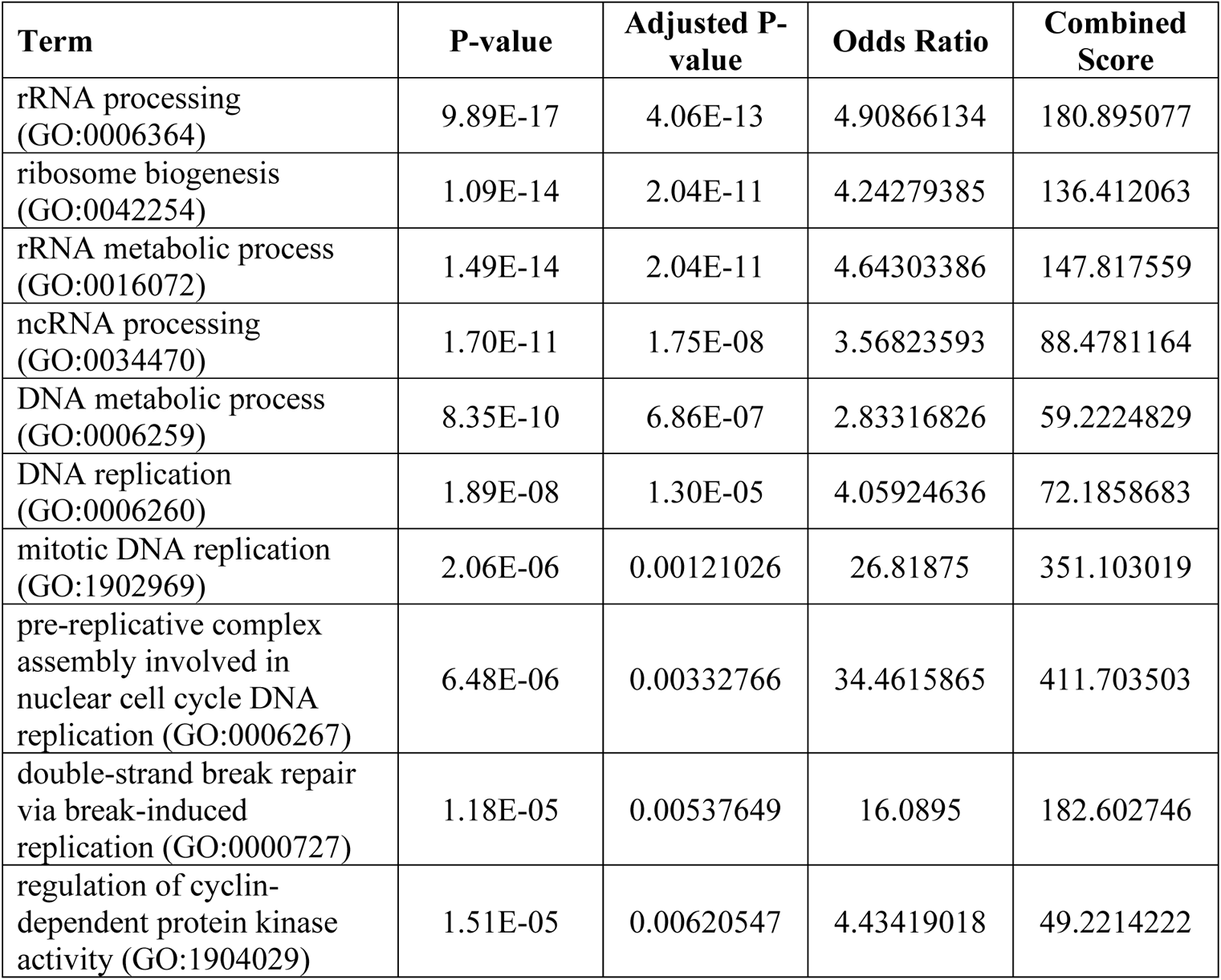
Biological processes associated with RAD21 knockdown. Top 10 GO Biological Processes for RAD21 DEGs sorted by adjusted p-value.

| **Term** | **P-value** | **Adjusted P-value** | **Odds Ratio** | **Combined Score** |
| --- | --- | --- | --- | --- |
| rRNA processing (GO:0006364) | 9.89E-17 | 4.06E-13 | 4.90866134 | 180.895077 |
| ribosome biogenesis (GO:0042254) | 1.09E-14 | 2.04E-11 | 4.24279385 | 136.412063 |
| rRNA metabolic process (GO:0016072) | 1.49E-14 | 2.04E-11 | 4.64303386 | 147.817559 |
| ncRNA processing (GO:0034470) | 1.70E-11 | 1.75E-08 | 3.56823593 | 88.4781164 |
| DNA metabolic process (GO:0006259) | 8.35E-10 | 6.86E-07 | 2.83316826 | 59.2224829 |
| DNA replication (GO:0006260) | 1.89E-08 | 1.30E-05 | 4.05924636 | 72.1858683 |
| mitotic DNA replication (GO:1902969) | 2.06E-06 | 0.00121026 | 26.81875 | 351.103019 |
| pre-replicative complex assembly involved in nuclear cell cycle DNA replication (GO:0006267) | 6.48E-06 | 0.00332766 | 34.4615865 | 411.703503 |
| double-strand break repair via break-induced replication (GO:0000727) | 1.18E-05 | 0.00537649 | 16.0895 | 182.602746 |
| regulation of cyclin-dependent protein kinase activity (GO:1904029) | 1.51E-05 | 0.00620547 | 4.43419018 | 49.2214222 |

**Table S3.** Biological processes associated with WAPL knockdown. Top 10 GO Biological Processes for WAPL DEGs sorted by adjusted p-value.

| **Term** | **P-value** | **Adjusted P-value** | **Odds Ratio** | **Combined Score** |
| --- | --- | --- | --- | --- |
| limb development (GO:0060173) | 6.87E-06 | 0.02950494 | 6.64022316 | 78.9432037 |
| lipid phosphorylation (GO:0046834) | 1.20E-04 | 0.19885464 | 14.0766234 | 127.106901 |
| cellular glucose homeostasis (GO:0001678) | 1.39E-04 | 0.19885464 | 4.07554745 | 36.1990471 |
| secondary alcohol biosynthetic process (GO:1902653) | 2.31E-04 | 0.24329865 | 4.49515399 | 37.6313707 |
| cholesterol biosynthetic process (GO:0006695) | 3.08E-04 | 0.24329865 | 4.30761719 | 34.8261574 |
| positive regulation of release of cytochrome c from mitochondria (GO:0090200) | 3.48E-04 | 0.24329865 | 5.28346124 | 42.0731483 |
| extracellular structure organization (GO:0043062) | 4.13E-04 | 0.24329865 | 1.95244613 | 15.2112593 |
| external encapsulating structure organization (GO:0045229) | 4.53E-04 | 0.24329865 | 1.94149067 | 14.9484369 |
| extracellular matrix organization (GO:0030198) | 6.16E-04 | 0.29117693 | 1.75673657 | 12.9851858 |
| sterol biosynthetic process (GO:0016126) | 6.78E-04 | 0.29117693 | 3.82835648 | 27.9342874 |

**Table S4.** NIPBL knockdown-associated biological processes rescued by co-depletion with WAPL. Top 10 GO Biological Processes for NIPBL DEGs rescued in the double knockdown condition sorted by adjusted p-value.

| **Term** | **P-value** | **Adjusted P-value** | **Odds Ratio** | **Combined Score** |
| --- | --- | --- | --- | --- |
| ribosome biogenesis (GO:0042254) | 2.79E-20 | 1.14E-16 | 5.277099 | 237.598868 |
| rRNA processing (GO:0006364) | 2.73E-18 | 5.58E-15 | 5.23727302 | 211.804312 |
| rRNA metabolic process (GO:0016072) | 9.23E-17 | 1.26E-13 | 5.13061397 | 189.429757 |
| ncRNA processing (GO:0034470) | 1.42E-14 | 1.45E-11 | 4.12911308 | 131.650349 |
| cytoplasmic translation (GO:0002181) | 3.79E-13 | 3.10E-10 | 6.15680228 | 176.092158 |
| SRP-dependent cotranslational protein targeting to membrane (GO:0006614) | 5.22E-12 | 3.56E-09 | 5.86102236 | 152.25747 |
| cotranslational protein targeting to membrane (GO:0006613) | 1.83E-11 | 1.01E-08 | 5.49351038 | 135.820725 |
| cellular macromolecule biosynthetic process (GO:0034645) | 1.98E-11 | 1.01E-08 | 2.91338179 | 71.8045298 |
| nuclear-transcribed mRNA catabolic process, nonsense-mediated decay (GO:0000184) | 1.33E-10 | 6.02E-08 | 4.63154902 | 105.336417 |
| ribosome biogenesis (GO:0042254) | 2.79E-20 | 1.14E-16 | 5.277099 | 237.598868 |

**Table S5.**
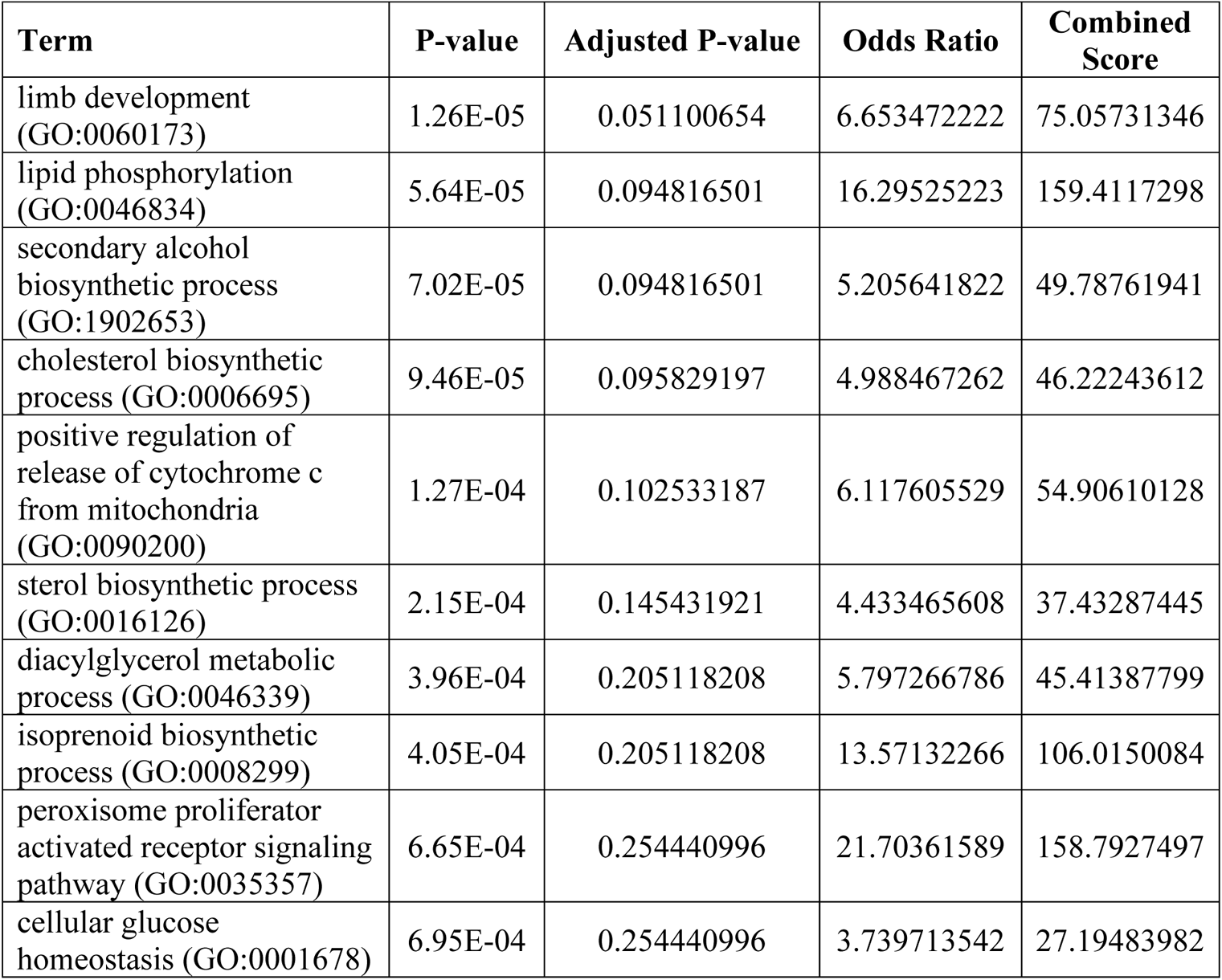
WAPL knockdown-associated biological processes rescued by co-depletion with NIPBL. Top 10 GO Biological Processes for WAPL DEGs rescued in the double knockdown condition sorted by adjusted p-value.

**Table S6.** Oligopaint design. Oligopaint design coordinates (hg19) and probe densities.

| **Probe** | **Chr** | **Start** | **Stop** | **# of Probes** | **Probes /kb** |
| --- | --- | --- | --- | --- | --- |
| Chr2:D1 | chr2 | 217531474 | 218621456 | 5464 | 5.01 |
| Chr2:D2 | chr2 | 218621456 | 220602102 | 8938 | 4.51 |
| Chr2:D3 | chr2 | 220602102 | 222889819 | 9479 | 4.14 |
| Chr2:S1 | chr2 | 219271354 | 219513415 | 765 | 3.16 |
| Chr2:S2 | chr2 | 219513415 | 219717782 | 846 | 4.14 |
| Chr2:S3 | chr2 | 219717782 | 219860895 | 656 | 4.58 |
| Chr2:S4 | chr2 | 219860895 | 220025303 | 939 | 5.71 |
| Chr2:S5 | chr2 | 220025303 | 220267857 | 1088 | 4.49 |
| Chr2:S6 | chr2 | 220267857 | 220406620 | 841 | 6.06 |
| Chr2:S7 | chr2 | 221552089 | 221965637 | 1640 | 3.97 |
| Chr2:S8 | chr2 | 221965637 | 222438280 | 2300 | 4.87 |
| Chr2:S9 | chr2 | 222438280 | 222889819 | 1635 | 3.62 |
| Chr3:S1 | chr3 | 45541203 | 45706813 | 678 | 4.09 |
| Chr3:S2 | chr3 | 45706813 | 45948482 | 1037 | 4.29 |
| Chr3:S3 | chr3 | 45948482 | 46120405 | 774 | 4.50 |
| Chr3:S5 | chr3 | 46705342 | 47040191 | 1763 | 5.27 |
| Chr3:S6 | chr3 | 47040191 | 47481841 | 1679 | 3.80 |
| Chr12:D1 | chr12 | 11707040 | 12890169 | 4676 | 3.95 |
| Chr12:D2 | chr12 | 12890169 | 13408925 | 2210 | 4.26 |
| Chr12:D3 | chr12 | 13408925 | 14338981 | 4211 | 4.53 |
| Chr19:D2 | chr19 | 17966474 | 18551439 | 2054 | 3.51 |
| Chr19:D3 | chr19 | 18551439 | 19097717 | 2502 | 4.58 |
| Chr19:S1 | chr19 | 17502026 | 17733289 | 679 | 2.94 |
| Chr19:S2 | chr19 | 17733289 | 17886271 | 514 | 3.36 |
| Chr22:D1 | chr22 | 33414401 | 35627637 | 7555 | 3.41 |
| Chr22:D2 | chr22 | 35627637 | 36520199 | 3271 | 3.66 |
| Chr22:D3 | chr22 | 36520199 | 36942436 | 1603 | 3.80 |
| MCM5 | chr22 | 35796115 | 35820495 | 120 | 4.92 |
| Chr22: part D2 | chr22 | 36019409 | 36520199 | 1673 | 3.34 |

**Table S7.** PRO-seq statistics. PRO-seq statistics, including raw read counts, mappable read counts to the spike in and reference genomes.

| **Samples** | **Number Raw Reads** | **Number Spike Reads Mapped** | **Number Unique Ref Reads Mapped** |
| --- | --- | --- | --- |
| PR28_control1 | 81763555 | 15317608 | 50020778 |
| PR30_control2 | 96354537 | 17005602 | 67235834 |
| PR32_Nipbl1 | 71081725 | 14234691 | 46820905 |
| PR34_Nipbl2 | 86026983 | 15549770 | 61607767 |
| PR33_Wapl1 | 90247992 | 18205253 | 51629855 |
| PR35_Wapl2 | 75712441 | 13925115 | 51202452 |
| PR29_NipblWapl1 | 88589172 | 14890161 | 61330222 |
| PR31_NipblWapl2 | 82623691 | 16999655 | 57261439 |
| PR36_NipblCtcf1 | 87588790 | 12047409 | 52428882 |
| PR38_NipblCtcf2 | 86547017 | 13443008 | 64122493 |
| PR37_Ctcf1 | 93071126 | 18664948 | 61286428 |
| PR39_Ctcf2 | 91546835 | 15866768 | 65933384 |

**Table S8.**
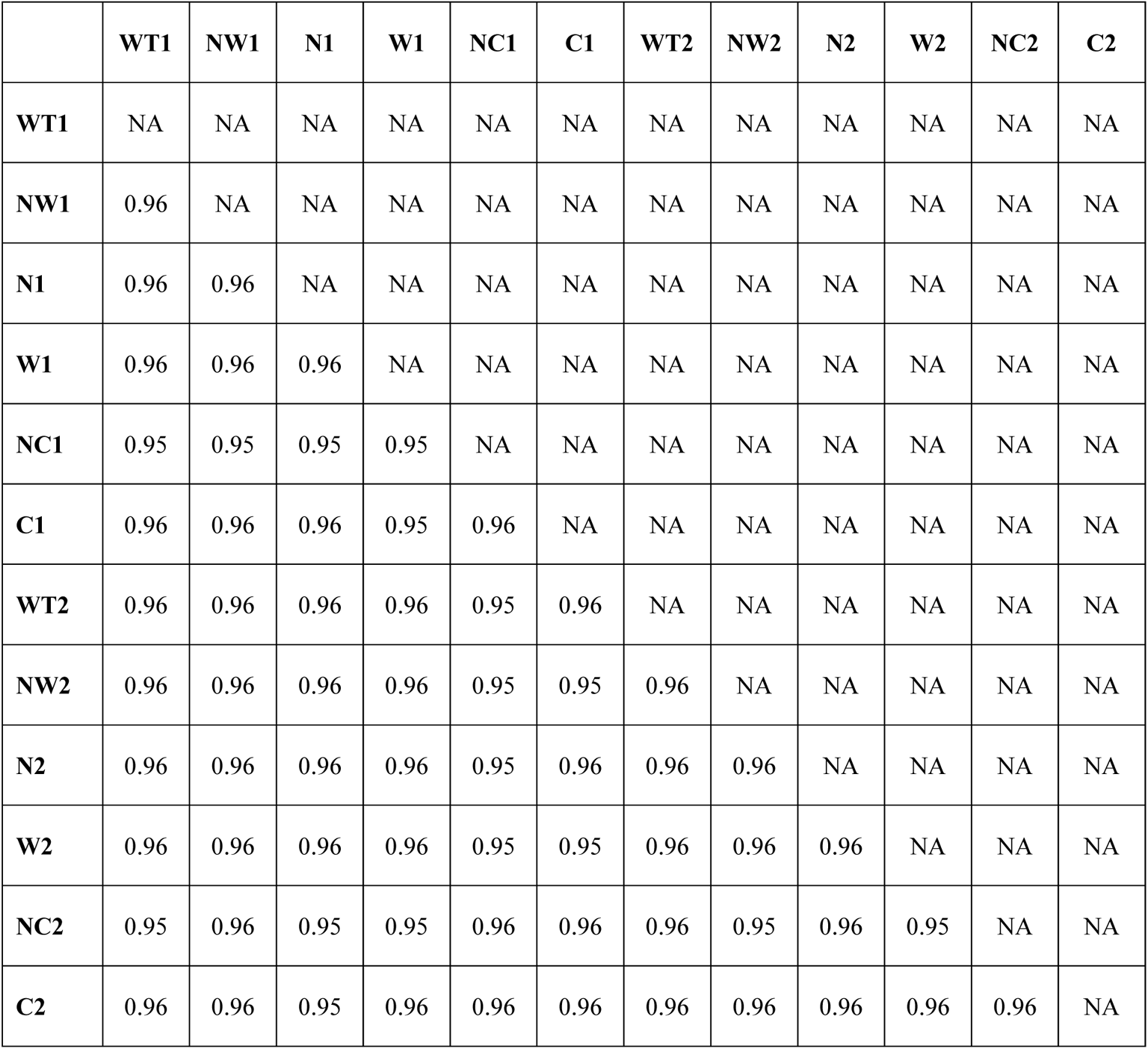
Pairwise correlation of PRO-seq counts. Pairwise correlation (Spearman’s rho) of PRO-seq counts in windows ±2kb around filtered TSS annotation. N represents NIPBL knockdown, W represents WAPL knockdown, C represents CTCF knockdown, NW represents NIPBL and WAPL double knockdown, and NC represents NIPBL and CTCF double knockdown.

